# Scalable imaging-free spatial genomics through computational reconstruction

**DOI:** 10.1101/2024.08.05.606465

**Authors:** Chenlei Hu, Mehdi Borji, Giovanni J. Marrero, Vipin Kumar, Jackson A. Weir, Sachin V. Kammula, Evan Z. Macosko, Fei Chen

## Abstract

Tissue organization arises from the coordinated molecular programs of cells. Spatial genomics maps cells and their molecular programs within the spatial context of tissues. However, current methods measure spatial information through imaging or direct registration, which often require specialized equipment and are limited in scale. Here, we developed an imaging-free spatial transcriptomics method that uses molecular diffusion patterns to computationally reconstruct spatial data. To do so, we utilize a simple experimental protocol on two dimensional barcode arrays to establish an interaction network between barcodes via molecular diffusion. Sequencing these interactions generates a high dimensional matrix of interactions between different spatial barcodes. Then, we perform dimensionality reduction to regenerate a two-dimensional manifold, which represents the spatial locations of the barcode arrays. Surprisingly, we found that the UMAP algorithm, with minimal modifications can faithfully successfully reconstruct the arrays. We demonstrated that this method is compatible with capture array based spatial transcriptomics/genomics methods, Slide-seq and Slide-tags, with high fidelity. We systematically explore the fidelity of the reconstruction through comparisons with experimentally derived ground truth data, and demonstrate that reconstruction generates high quality spatial genomics data. We also scaled this technique to reconstruct high-resolution spatial information over areas up to 1.2 centimeters. This computational reconstruction method effectively converts spatial genomics measurements to molecular biology, enabling spatial transcriptomics with high accessibility, and scalability.

## Introduction

Tissue functions arise from the coordinated activities of cells. Interactions at multiple scales – from the level of molecules to communication between cells, to signaling across tissues – are all necessary for this coordinated function. Spatial transcriptomic technologies, which enable spatial localization of gene expression profiles within tissue contexts, represent a powerful set of tools to study tissue function and cell-cell interactions^1–5^. Currently, imaging plays a key role in spatial transcriptomics, either for directly locating RNA molecules within tissues or for indexing capture arrays. However, imaging, which requires specialized techniques and equipment, introduces several limitations on spatial transcriptomic approaches, such as throughput, adaptability and the constrained size of detectable areas^6^. Alternatively, arrays may be deterministically printed through lithography or physical methods, but such methods require complex equipment and high upfront costs^3,7^. An imaging-free spatial transcriptomic technique, which ideally can be performed without complex equipment, can enhance the throughput and accessibility of experiments, and enable larger scale detection for comprehensive studies of tissues.

In theory, spatial information can be inferred independently of imaging. For example, molecular proximity measurements measure interactions (e.g. HiC^8^). Beyond proximity, spatial locations can be inferred directly from pairwise distance measurements. For example, the location of a mobile phone is accurately determined by measuring its distances to three different satellites. Similarly, the geographical layout of the United States can be mathematically reconstructed based solely on the pairwise distances between cities^9^. Moreover, the concept of distance can be generalized from euclidean distance to more complicated spatial variation. Human genetic variations, for example, tend to correlate geographically; the two-dimensional representation of European genetic variations has been shown to mimic the actual geography of Europe, revealing the geographic information that are inherent within genetic variations^10^. At the molecular level, diffusion patterns, which highlight neighboring information, have been used for reconstructing the spatial locations of molecules^11–13^. These efforts to determine molecular locations without traditional imaging have broadened our methodologies for conducting spatial measurements. However, current approaches are either established only in theoretical simulations^14,15^, or in simplified experimental systems^13^.

Here, we developed an imaging-free spatial transcriptomics method that computationally reconstructs the spatial locations of barcode arrays used in spatial transcriptomics measurements with high resolution and fidelity. We implemented the imaging-free approach on two-dimensional barcode arrays, along with ground truth imaging for error estimation. Then we utilized a dimensionality reduction method to reconstruct the spatial locations. We demonstrated that this imaging-free strategy can be integrated with existing barcode array based spatial transcriptomics methods without perturbing spatial structures. This method facilitates higher throughput generation of barcode arrays and is accessible to laboratories lacking specialized imaging equipment. Furthermore, we applied this technique to a tissue sample on a centimeter scale, showing its potential for large-scale spatial transcriptomics.

## Results

### Computational reconstruction of spatial locations through dimensionality reduction

We started by testing whether computational reconstruction can map spatial locations using diffusion based proximity data. To do so, we implemented a framework to generate simulated diffusion data. Since Slide-seq, a high resolution spatial transcriptomic approach, utilizes arrays of barcoded beads for spatial capture, we theorized that we could simulate diffusional interactions between beads on a 2D array (Fig.1a). To visualize the reconstruction effect, we uniformly sampled capture and fiducial beads from a circular area with color pattern of a letter ‘H’ (Fig.1b, Supplementary Fig.1a). The diffusion of barcodes from each fiducial bead was simulated to follow a Gaussian distribution (Supplementary Fig.1b, Methods). Due to this diffusion, capture beads exhibit proximity-dependent capture of barcodes from fiducial beads, generating a neighboring matrix between bead barcodes (Fig.1a). Specifically, capture beads registered higher count values for barcodes from fiducial beads that were closer, while distant fiducial beads were associated with zero counts. While each capture barcode can be characterized by count values on its associated fiducial barcodes, physically adjacent capture beads capture similar barcodes from similar fiducial beads and are closer in the high-dimensional space of fiducial bead barcodes. We reasoned that dimensionality reduction algorithms may be able to reconstruct the latent two-dimensional representation of physical space from this high-dimensional interaction matrix. In particular, we performed Uniform Manifold Approximation and Projection^16^ (UMAP), a nonlinear dimensionality reduction method, which reduces the high-dimensional capture bead barcode by fiducial bead barcode data into a two-dimensional embedding space, while preserving the similarity of capture bead barcodes in high dimensional space (Supplementary Figure 1). While all parameters clustered by proximity, we found a range of parameters (Supplementary Figure 2) wherein locations of capture beads in this two-dimensional embedding are highly similar to their physical locations, indicating UMAP learns the intrinsic two-dimensional manifold embedded within the high-dimensional data (Fig.1b, Supplementary Fig.1c,d). The accuracy of reconstruction was quantitatively assessed by comparing the reconstructed locations of the capture beads to their known physical positions, following a rigid transformation for alignment. The median error in simulation was found to be equivalent to 1.6% of the array’s diameter (Supplementary Fig.1e,f,g, Supplementary Fig.2b). Other dimensionality reduction methods, encompassing both linear and nonlinear, were assessed using identical simulation data (Supplementary Fig.3). While all methods showed promise in discerning the order of bead arrangement, they incurred larger absolute errors when compared to the results of UMAP.

**Figure 1:**
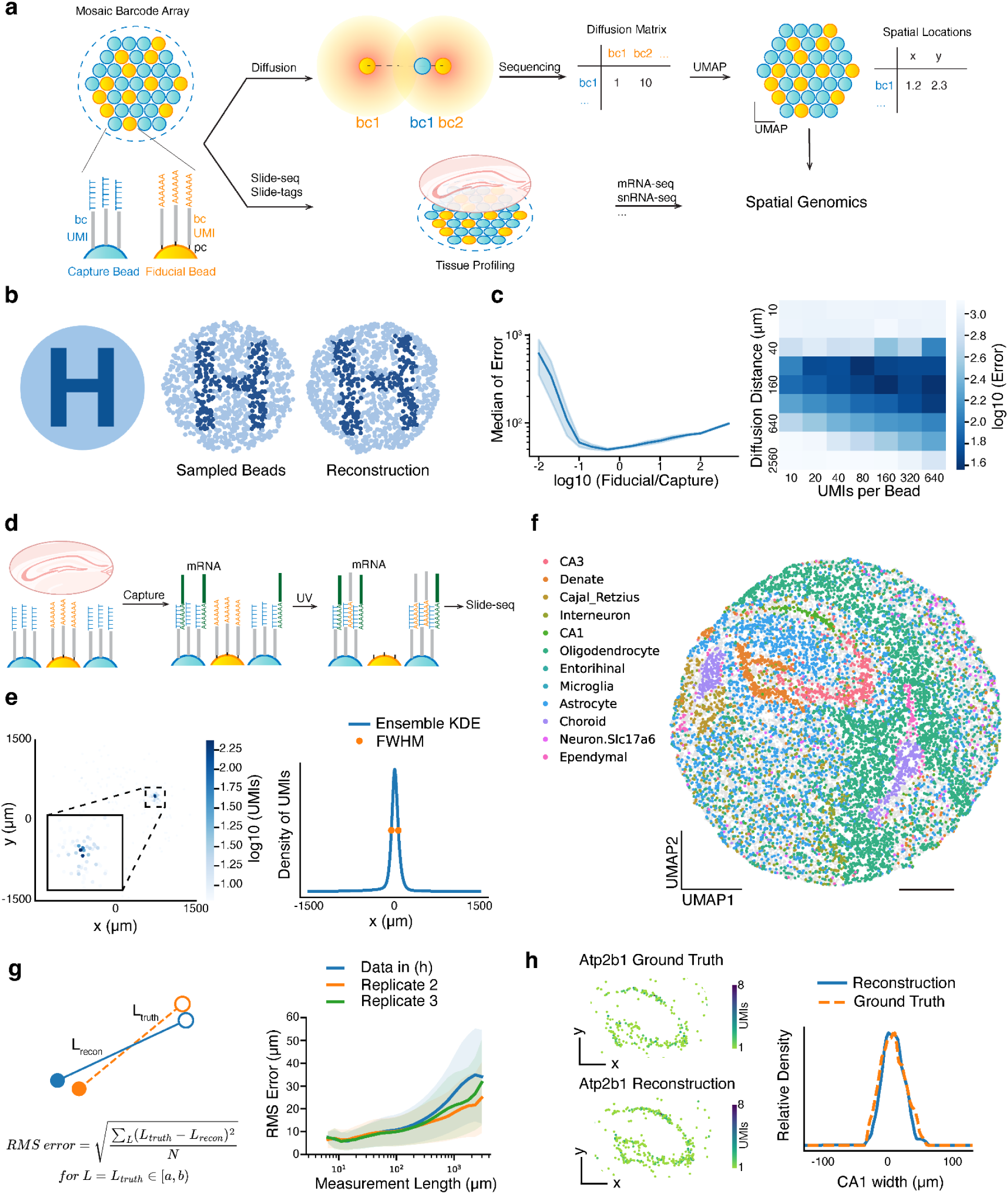
Imaging-free spatial transcriptomics through computational reconstruction. **a**, Schematic of imaging-free reconstruction for spatial transcriptomics. A mosaic barcode array of uniformly mixed capture beads (poly(dT)) and fiducial beads (poly(dA)) is used for imaging-free spatial transcriptomics. Both capture and fiducial beads are DNA-barcoded with a unique spatial barcode for each bead and a unique molecular identifier (UMI) for each oligonucleotide molecule. Fiducial beads’ DNA barcodes are photocleaved and diffuse to proximate capture beads. A diffusion matrix can be abstracted from the sequencing result of capture bead barcode and fiducial bead barcode conjugation product. UMAP reduces the high dimensional diffusion matrix to a two dimensional embedding space and reconstructs the spatial location of beads. With computationally reconstructed spatial locations, the same array is performed with Slide-seq/Slide-tags to profile tissue spatial genomics. bc, spatial barcode. pc, photocleavable linker. **b**, Simulated reconstruction. (Left) An image with the pattern of letter ‘H’. (Middle) Beads sampled from the original image. (Right) Reconstructed beads location through simulated diffusion matrix and UMAP embedding. Each bead is colored the same as the middle image to show the recovered pattern. **c**, (Left) Simulated reconstruction error as a function of the ratio of fiducial to capture beads. The error is quantified as the median displacement distance of capture beads, with the whole array’s diameter defined as 3 mm. The shaded area represents one standard deviation (N = 5 repeated simulations). (Right) Simulated reconstruction error as a function of diffusion distance and number of molecules carrying diffusion information (UMIs) per bead. **d**, Schematic of reconstruction for Slide-seq. mRNA molecules are captured by capture beads, followed by UV induced fiducial bead barcode diffusion and conjugation. **e**, (Left) Diffusion pattern of a capture bead barcode on its associated fiducial bead barcodes, colored by the number of unique joint UMIs. Inset detailing the diffusion center. (Right) Ensemble average of kernel density estimation (KDE) on capture bead barcode’s diffusion distribution in x-axis direction. Orange dots represent the half-maximum, corresponding to the full width at half maximum (FWHM) measured as 123.1 µm. **f**, Spatial location of capture beads from reconstruction through UMAP embedding, colored by decomposed cell types. Locations were scaled to the original array size with a diameter of 3 mm. **g**, (Left) Schematic of RMS error at different measurement lengths. Error is defined as the difference between the distance of two beads measured in ground truth (orange dashed line) and in reconstruction (blue line). Errors of measurement lengths in the same range are averaged by root-mean-square (RMS). (Right) RMS error of different measurement lengths. Data shown in (f) (blue) and two biological replicates (orange and green) are presented. Solid lines represent mean values across beads and shaded areas represent one standard deviation. **h**, (Left) Spatial expression of Atp2b1, a CA1 layer marker, in ground truth and reconstruction. (Right) Representative plots of CA1 layer width by profiling the expression intensity of Atp2b1 along a perpendicular line in ground truth (orange) and reconstruction (blue). Scale bars: 500 µm.

To assess the practical applicability of this computational reconstruction method in experiments, we explored this method’s dynamic range by tuning various parameters in simulation. We analyzed reconstruction errors with different ratios of fiducial to capture bead numbers. With the ratio ranging from 1:5 to 5:1, the simulation displayed median errors less than 2% of the array size (Fig.1c). This range of ratio was subsequently considered for the design of experimental mixed arrays. We also simulated the effects of changing captured unique molecular identifiers (UMIs) per capture bead and diffusion distance (σ) (Fig.1c). Both excessively narrow and wide diffusion distance led to larger errors, whereas a greater number of UMIs per bead enabled better positioning accuracy. We were encouraged that a wide range of parameters demonstrated feasibility for computational reconstruction, for example diffusion distance (σ) between 2% to 6% of the array size, and a minimum of 40 UMIs per bead.

### Implementing spatial transcriptomics through computational reconstruction

We next sought to implement and validate our reconstruction strategy experimentally by performing a Slide-seq assay on an array that was both spatially indexed and reconstructed by our new approach. Our array mixed the original barcoded poly(dT) beads for capturing mRNA with barcoded poly(dA) fiducial beads to enable diffusion-based reconstruction (Fig.1a,d). Next, we performed in situ sequencing to spatially index the array using our standard approach, as previously described^17^, to generate ground truth positions before reconstruction. To obtain spatial coordinates of beads by reconstruction, the oligonucleotide barcodes on poly(dA) beads (fiducial beads) were cleaved with UV light, enabling diffusion to nearby poly(dT) beads (capture beads) (Fig.1d). The distribution of capture bead barcodes on fiducial bead barcodes followed a heavy tailed distribution, with the full width at half maximum (FWHM) around 123.1 µm (Fig.1e).

Given that this approach generates diffusion of spatial information from fiducial beads to capture beads, we next sought to reconstruct the relative spatial locations. We applied UMAP on the high-dimensional diffusion information to reconstruct the relative locations of capture beads in two-dimensional embedding space, without any spatial information input. In addition to the two main UMAP parameters we tuned in simulation, we also found increasing the number of epochs, using cosine metric for computing high dimensional distance, and applying log1p transformation of diffusion matrix improved the reconstruction accuracy (Supplementary Fig.5). Comparing to ground truth from in situ sequencing, reconstructed locations recovered the global arrangement of capture beads (Supplementary Fig.5a). We noticed a few beads (21 out of ~16000) were positioned significantly away (>200 µm) from ground truth positions, most of which were attributable to in situ sequencing errors (Supplementary Fig.6) or were barcode collisions.

Given the reconstructed bead locations, we next examined spatial transcriptomics data captured by the array. We performed Slide-seqV2 using the same reconstructed array with the capture beads (poly(dT)) on a mouse hippocampal transcript with no modifications to the protocol. Individual bead profiles were clustered and assigned with cell types by robust cell type decomposition^18^ (RCTD) (Supplementary Fig.4d). Spatial representation of cell types with reconstructed capture bead location demonstrated the known structures of the hippocampus (Fig.1f, Supplementary Fig.4d). This is especially clear when examining cell-type distributions and marker gene plots, which are virtually indistinguishable between reconstruction and ground truth data (Supplementary Fig.7, 8).

We next sought to quantify the accuracy of the reconstruction process. To do so, reconstruction accuracy was assessed with three strategies: examining the absolute error, relative error, and histological structure preservation in comparison to ground truth. First, to assess absolute error, we calculated each capture bead’s absolute displacement via a rigid registration between reconstruction result and ground truth. The median value of displacement lengths was 25.9 µm (Supplementary Fig.4a,b,c). The absolute error can be affected by the registration process; this suggests that a more appropriate statistic should be the error in distance measurements between reconstruction and ground truth (Fig.1g, Supplementary Fig.4e). The intuition here is that most commonly, we are quantifying pairwise distance measurements (e.g. length, neighborhoods, spatial proximity), and thus, we care about the error of pairwise distance measurements. We quantified the root-mean-square (RMS) error of length measurements across all pairwise length measurements as a function of measurement length. RMS error was close to 10 µm (the bead size) at local scale measurements (~100 µm) as nearby beads were usually displaced in the same direction, and plateaus at ~25 µm >1000 micron (representing <2.5% error in measurement lengths) (Supplementary Fig.4e). To assess the effect of reconstruction error on histological structure, we measured the width of the CA1 layer in the hippocampus. The widths, characterized by CA1 marker Atp2b1 expression, were similar in ground truth and reconstruction (FWHM is 49.5 µm in ground truth and 43.7 µm in reconstruction) (Fig.1h). We also performed neighborhood enrichment analysis (Methods) between all pairs of cell types and found the results highly similar between reconstruction and ground truth (with Pearson correlation coefficient = 0.997) (Supplementary Fig.4g). Lastly, we evaluated the reconstruction error across three biological replicates in the hippocampus, and found that results were largely concordant across replicates, thus demonstrating the robustness of the reconstruction procedure (Fig.1g, Supplementary Fig.4f). We also found that across the three replicates, the gene expression information with matched spatial barcode increased by 1.6 times in reconstruction compared to ground truth (Supplementary Fig.4h). These data demonstrate that the reconstruction error is relatively small (~25 µm, 2.5 times of bead size), with a subtle effect on local spatial transcriptomics analysis.

### Spatial reconstruction at single-nucleus resolution

As the computational reconstruction strategy should be adaptable to all array-based spatial technologies, we next sought to apply reconstruction with Slide-tags, a recently developed single-nucleus spatial technology^19^.

In Slide-tags, nucleic acid spatial barcodes are photocleaved from barcoded arrays, associated with nuclei, followed by single-nucleus RNA sequencing with single-cell indexing. To apply reconstruction to Slide-tags, we generated arrays with mixtures of photocleavable poly(dA) (fiducial) and non-cleavable poly(dT) (capture) beads. In the Slide-tags experiment, the fiducial beads (poly(dA)) are photocleaved, diffused, and captured by capture beads while tagging nuclei at the same time (Fig.2a).

**Figure 2:**
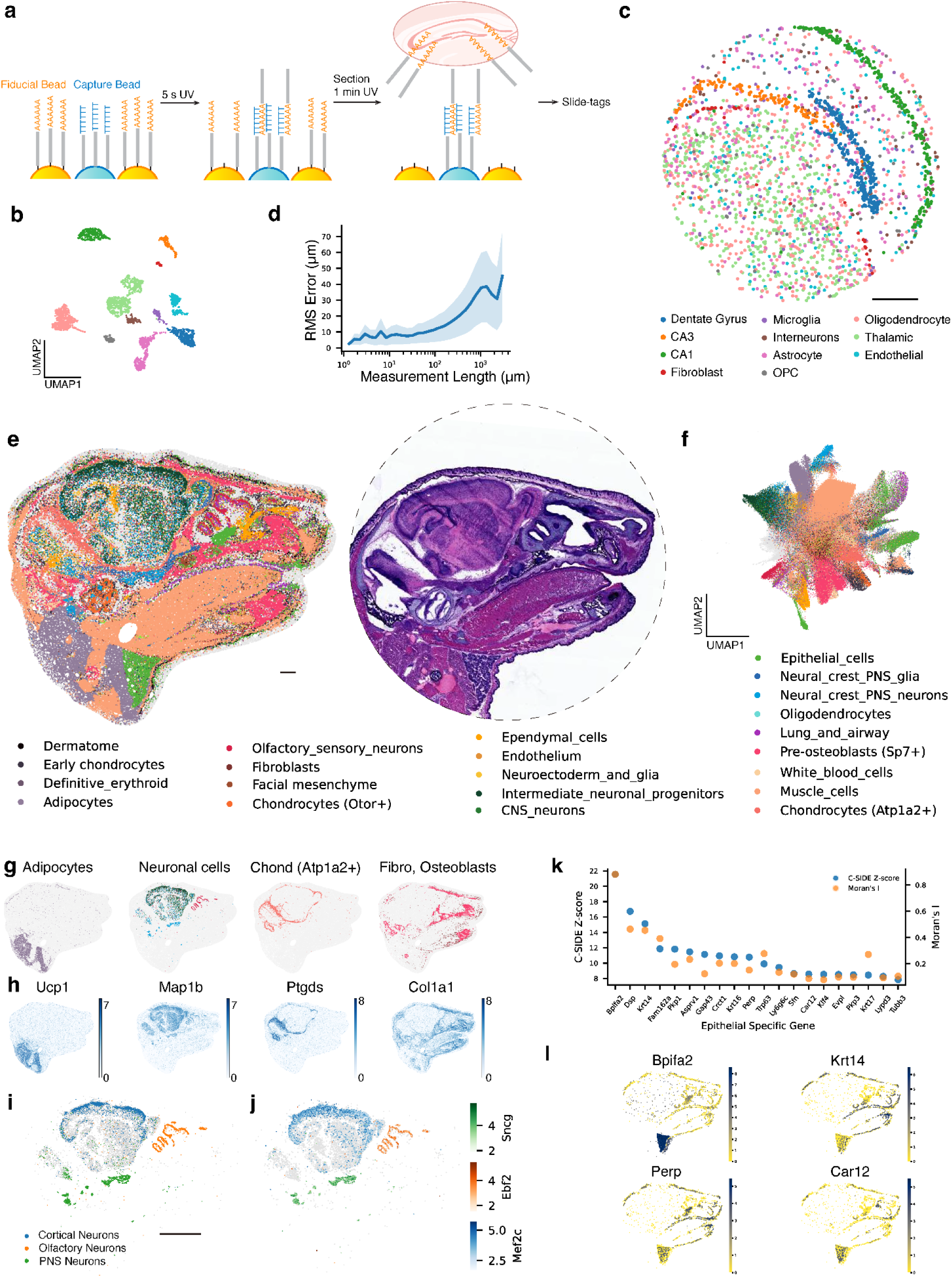
Reconstruction enables diverse spatial transcriptomics measurements at scale. **a**, Schematic of reconstruction for spatial transcriptomics with single-nucleus resolution. A barcode array is exposed to UV for a 5 s to allow fiducial bead barcode diffusion and conjugation with capture bead barcode. After the tissue section, the array is UV exposed for 1 min to cleave majority of fiducial bead barcode and tag nuclei. Sample is to proceed with Slide-tags for spatial genomics profiling. **b**, UMAP embedding of snRNA-seq profiles from a coronal mouse hippocampus section, colored by cell type annotations shown in c. **c**, Spatial location of nuclei mapped based on reconstructed locations of capture beads, colored by cell type annotations. Locations were scaled to the original array size with a diameter of 3 mm. **d**, RMS length measurement error of nuclei in reconstruction versus ground truth. Solid line as the mean value and shaded area as one standard deviation. **e**, (Left) Spatial location of beads with profiled gene expression from a P1 mouse head section. Beads are colored by decomposed cell type annotations from RCTD. (Right) Hematoxylin and eosin staining of an adjacent slice from the same sample. Dashed line indicates the 1.2 cm bead array used for this experiment. **f**, UMAP representing gene expression of the mouse sample captured by beads, colored by decomposed cell types from RCTD. **g**, Spatial distribution of labeled cell types, colored the same as cell type annotations in f. Gray beads represent other cell types, plotted for contrasting. Neuronal cells include olfactory sensory neurons, intermediate neuronal progenitors, CNS neurons, and neural crest PNS neurons from clusters in e. Chond, chondrocytes. Fibro, fibroblasts. **h**, Marker gene spatial expression of cell types shown in h. Colored by relative expression level. **i**, Subclustring of neuronal cells, colored by subtype annotations. Gary beads are other subclusters of neuronal cells. **j**, Maker gene of each neuronal subtype shown in i, colored with corresponding color gradients based on relative expression level. All beads of neuronal cell type are plotted. **k**, Within beads categorized under the epithelial type, top 20 spatially differential expression genes ranked by nonparametric C-SIDE were plotted, with Moran’s I statistics calculated. Higher scores on both metrics signify more spatially variable expression. **l**, Spatially differential expression of four epithelial genes from k are shown. All beads of epithelial type are positioned, colored by relative expression level of each gene. Scale bars: 500 µm.

We performed reconstruction Slide-tags on a mouse hippocampal section. Again, in addition to computational reconstruction of the array, we performed in situ sequencing to generate ground truth spatial positions on the same array. We generated high-quality single-nucleus spatial data (2091 genes per cell), which shows high quality clustering of major cell types in the hippocampus (Fig.2b, Supplementary Fig.10). We next reconstructed the bead locations with diffusion information and positioned each nuclei based on bead barcodes that it is tagged with. Following reconstruction and spatial placement, the spatial representation of different cell types captured the architecture of the hippocampus (Fig.2c).

We next evaluated the accuracy of the reconstruction of Slide-tags. We calculated the absolute positioning errors of each bead and each nucleus by comparing to ground truth locations from in situ sequencing, after applying a rigid transformation (Supplementary Fig.9). We found that the median value of bead positioning error (25.4 µm) and nuclei positioning error (27.2 µm) are similar (Supplementary Fig.9). To avoid the error introduced by registration, we also calculated the the RMS error of measurement lengths, which was smaller than 25 µm at local measurement scales (<500 µm) and plateaus at ~30 µm >1000 microns (representing <3% error in measurement lengths) (Fig.2d, Supplementary Fig.9g). To assess the effect of reconstruction error on biological structure, we compared the spatial representation of each clustered cell type in reconstruction and ground truth. We found no detectable difference in the dimensions of brain structures (Supplementary Fig.11). Lastly, Slide-seq and Slide-tags showed similar performance with respect to reconstruction error, likely due to the shared characteristics of the reconstruction protocol.

### Computational reconstruction enables spatial transcriptomics at large scale

Our reconstruction technique is purely performed through molecular biology reactions, and, thus, is not limited by imaging throughput. To demonstrate the scalability of reconstruction for spatial transcriptomics, we performed reconstruction on a 1.2 centimeter P1 mouse cranial section with Slide-seq (Supplemental Video 1). We spatially profiled transcriptomics across different tissue types, including brain, muscle, and the upper respiratory system, with a single section (Fig.2e). When compared with hematoxylin and eosin staining of an adjacent section, reconstruction successfully identified the compartmentalization of different tissues and elucidated fine structural details (Fig.2e). We assigned decomposed cell types to each bead with RCTD (Fig.2f, Supplementary Fig.12c, Supplementary Fig.13). To assess the fidelity of reconstructed tissue structure, we represented the spatial distribution of certain cell types that exhibit unique spatial localization: adipocytes around anterior cervical region; neuronal cells in central nervous systems (CNS), peripheral nervous systems (PNS), and olfactory sensory region (Fig.2g). The locations of these cell types are highly correlated with the spatial expression pattern of their maker genes (Fig.2h). For instance, the locations of fibroblasts and osteoblasts correspond with the distribution of type I collagen, which is abundantly present in tendons and bones. To examine the spatial transcriptomics data in detail, we gathered beads that were assigned with neuronal cell types and subjected them to further clustering. Such subclustering revealed distinctions between CNS, PNS, olfactory neurons, which were highly correlated with the expression pattern of their respective marker genes. We found cortical neurons with this higher resolution of clustering (Fig.2i,j, Supplementary Fig.12d,e,f).

With this comprehensive spatial transcriptomics profiling, we sought to identify genes with spatially differential expressions. We first focused on the olfactory epithelium for its well-organized structures and multilayered cell compositions. We profiled the spatial expression of previously identified genes and found the result consistent with in situ hybridization^20^: carbonyl reductase 2 (*Cbr2*), a sustentacular cell maker, has higher expression in olfactory epithelium’s outer layer; regenerating islet-derived protein 3 gamma (*Reg3g*), a respiratory epithelium marker, has a significant expression decrease at the boundary between respiratory epithelium and olfactory epithelium; and growth associated protein 43 (*Gap43*), which marks immature olfactory sensory neurons (OSNs), is only expressed in the inner olfactory epithelium layer where immature OSNs are located (Supplementary Fig.14a). Furthermore, by analyzing genes enriched in the olfactory epithelium, we profiled additional spatially variable genes in the olfactory epithelium (Supplementary Fig.14b). The spatial profiling of spatially differential expression genes in olfactory epithelium further displayed the high fidelity of our reconstruction method at fine structural details.

As our data set covers a wide range of tissue types, we sought out to identify cell type-specific differential gene expression across the entire section. We performed nonparametric cell type-specific inference of differential expression^21^ (C-SIDE) of epithelial specific genes to identify genes with spatially variable expression. We also calculated Moran’s I statistics, a spatial autocorrelation measurement, of genes with high C-SIDE Z-scores. The results from both analyses indicate a nonrandom pattern in the variable expression of these genes (Fig.2k). Spatial representation of selected genes displayed their distinct expression patterns across various regions (Fig.2l). For example, BPI fold containing family A member 2 (BPIFA2) is prominently expressed in the parotid gland epithelial cells. In addition, we analyzed spatial differential expression in muscle cells, in a similar manner, and found 277 region specific muscle genes (Supplementary Fig.15, Supplemental Table 1). These analyses demonstrate our reconstruction method is capable of discovering spatially relevant genes and revealing the complexity of gene expression patterns.

## Discussion

Our work illustrates that a molecular diffusion based computational reconstruction approach can enable imaging-free spatial transcriptomics with high fidelity. In particular, we demonstrated that UMAP can reconstruct a two dimensional barcode array given a diffusive interaction process between barcodes. In this process, UMAP successfully learns the intrinsic two dimensional manifold embedded within the high-dimensional diffusion data. The rationale of using a dimensionality reduction method for this reconstruction task is that the bead barcode diffusion process generates information in a high dimensional space in which physically closer bead barcodes are also closer. Thus, to reconstruct the physical locations, a method should preserve the high dimensional distances while reducing it to a low dimensional space, which matches the aim of dimensionality reduction methods. Intuitively, nonlinear dimensionality reduction methods may be preferred due to the nonlinear nature of the diffusion process. Furthermore, in contrast to t-Distributed Stochastic Neighbor Embedding (t-SNE), which primarily conserves local distances, UMAP is adept at preserving a broader spectrum of distances^22^. Empirically, we found that UMAP outperforms a variety of other dimensionality reduction algorithms with the highest accuracy and the shortest running time.

We systematically assessed the error of reconstruction. The assessment of absolute error, by direct comparison between the actual and reconstructed bead locations may overestimate errors, as reconstruction deformation may be locally smooth. Therefore, evaluating the relative error in distances between pairs of beads provides a more reliable metric. We noticed that within the range of several hundred micrometers, bead displacements tend to be concordant. Thus, for small measurement lengths (<500 µm), the error remains small, typically under 20 µm. Lastly, we evaluated the distortion of reconstruction when analyzing biological structure such as the mouse hippocampus and showed that such locally concordant errors have a minor effect on local structures or neighboring cell analyses. Thus, computational reconstruction through dimensionality reduction generates high fidelity, and high-resolution spatial transcriptomics data.

Despite the high fidelity of the reconstruction strategy given its simplicity, it nonetheless introduces some variation into the data. Higher accuracy in reconstruction could potentially be attained by solving it as an assignment problem with predefined locations; for instance,by imaging the bead array in bright field conditions and identifying the locations of beads while the corresponding barcodes remain unidentified. Lastly, while reconstruction decouples the scale of spatial transcriptomics from imaging, the sequencing cost of reconstruction grows linearly with the array area. While costs are currently modest for reconstruction (Supplemental Table 2), we anticipate they will continue to decrease with the exponential decrease in sequencing costs and adoption of novel sequencing technologies.

This work demonstrates that computational reconstruction can enable spatial transcriptomics at a large scale and high throughput. An elegant aspect of molecular biology tools supporting modern day genomics (e.g. RNA-seq, epigenomics) has been the ability to distribute through open-source protocols and readily available enzymes. Here, we demonstrate that we can effectively convert spatial transcriptomics into a molecular biology tool, as opposed to one that requires specialized equipment, thus, allowing for widespread accessibility for the scientific community. While we currently demonstrate the technology with Slide-seq and Slide-tags, the experimental and computational approach may be highly generalizable to array-based capture technologies, as well as potentially directly within tissues in 3D contexts. Lastly, decoupling from microscopy will enable us to perform spatial genomics at spatial scales not limited by microscopy, to reach the dimensions of entire human organs.

## Supplementary Figures

**Supplemental Figure 1:**
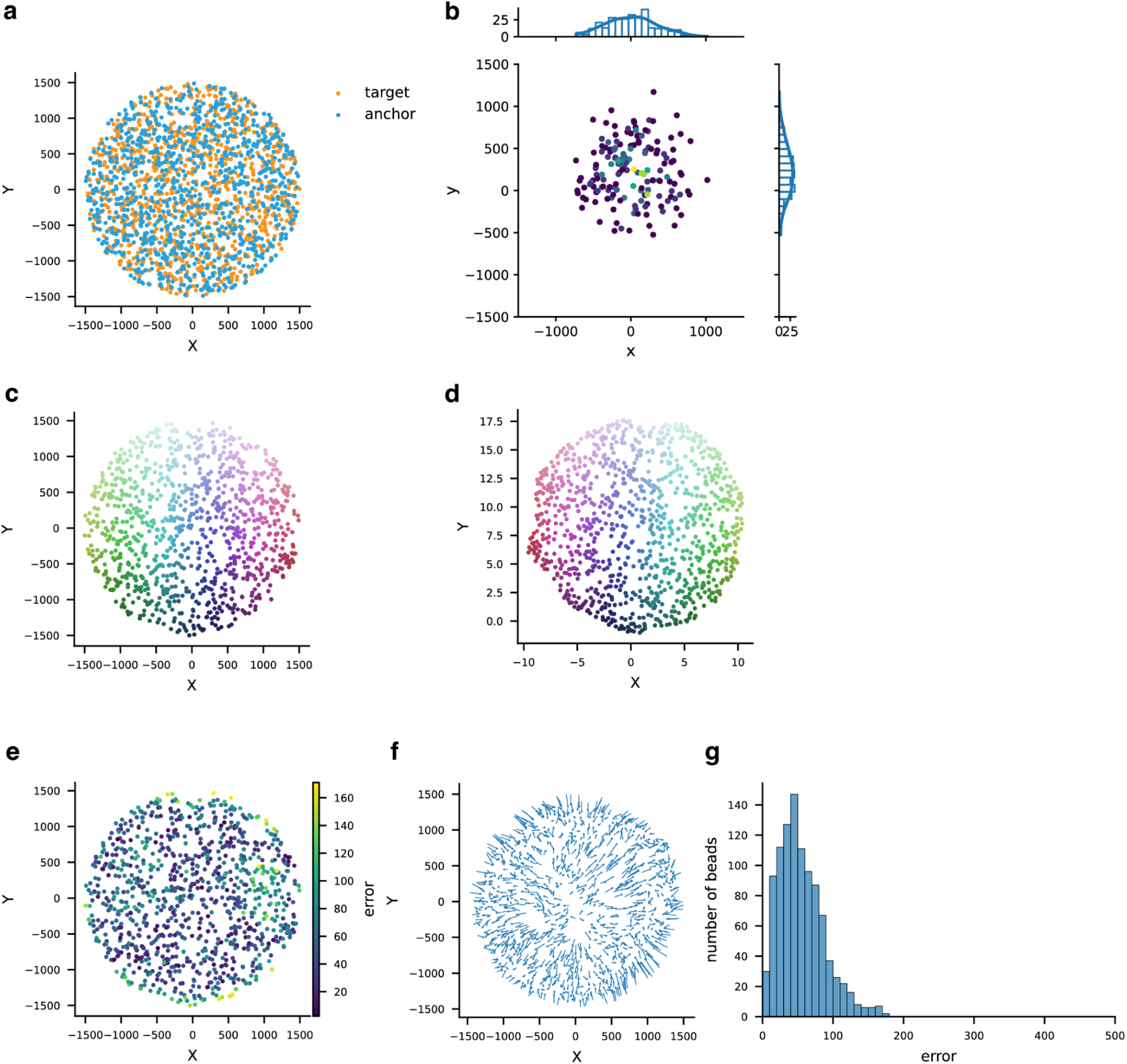
Simulation of diffusion and reconstruction with UMAP. **a**, Simulated locations of capture beads and fiducial beads in a 3 mm circle. **b**, Simulated diffusion pattern of a capture bead on its associated fiducial beads, colored by simulated UMI counts. The distribution plots on the top and right represents the diffusion distribution on the x and y axis respectively. **c**, Simulated locations of capture beads, colored by a two dimensional color gradient depending on the locations. **d**, UMAP reconstructed locations of capture beads, colored the same as in c. **e**, Absolute error of capture beads plotted in ground truth locations. **f**, Displacement vectors of capture beads. Each arrow starts from the capture bead’s ground truth location and ends at the reconstruction location. **g**, Histogram plot of capture beads’ absolute error.

**Supplemental Figure 2:**
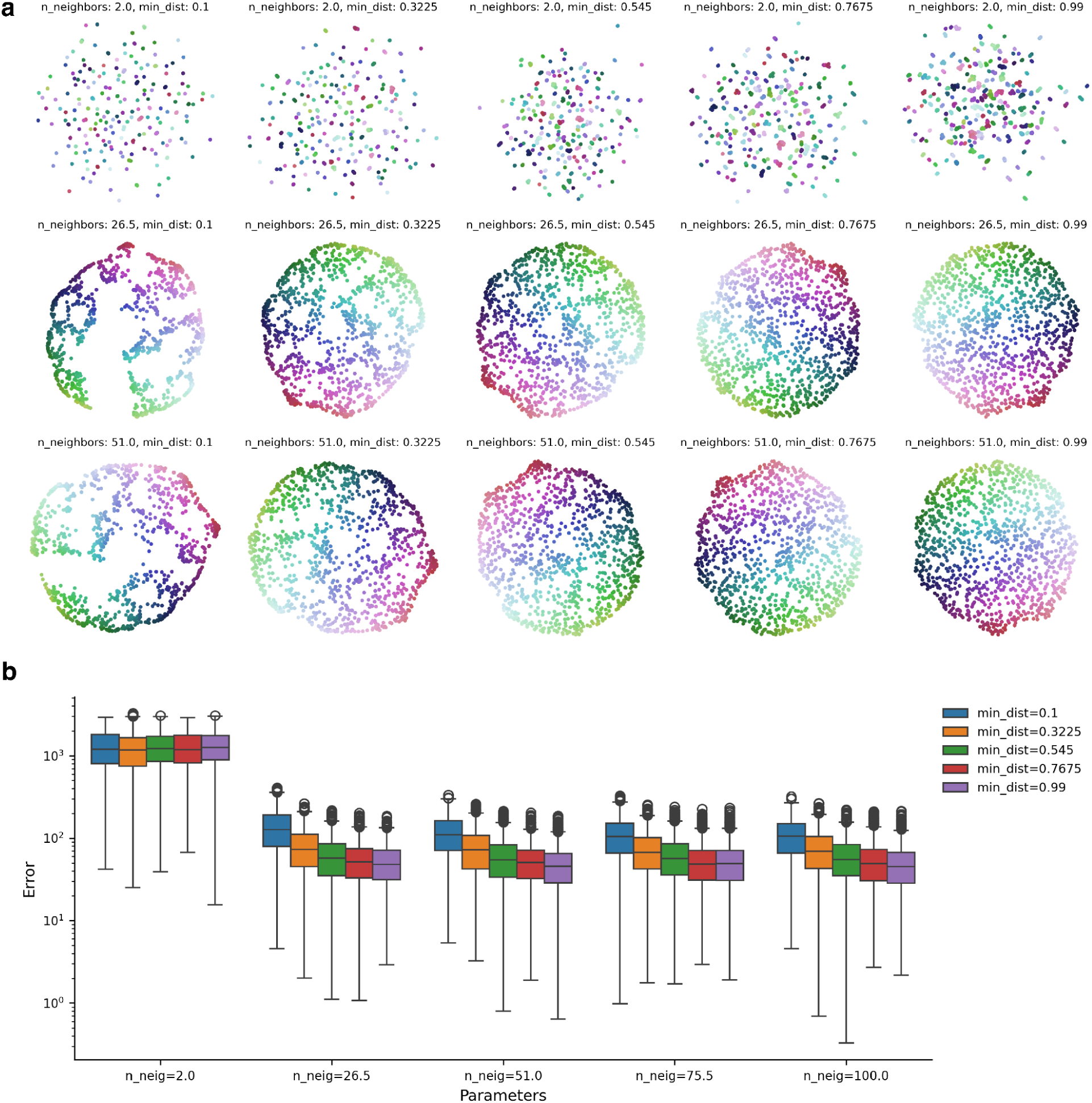
UMAP parameters’ effect on reconstruction. **a**, The same diffusion matrix is embedded in the two dimensional space with different parameters of UMAP: n_neighbors as the size of local neighborhood and min_dist as effective minimum distance between embedded points^16^. Each bead is colored the same as in Supplemental Figure 1c to show the pattern recovery. **b**, Boxplot of reconstruction error with different UMAP parameters. Each box represents absolute errors of beads with corresponding parameters, with middle box as error between 25% to 75%, middle line as median error, and outlier as 1.5 times of interquartile range (IQR).

**Supplemental Figure 3:**
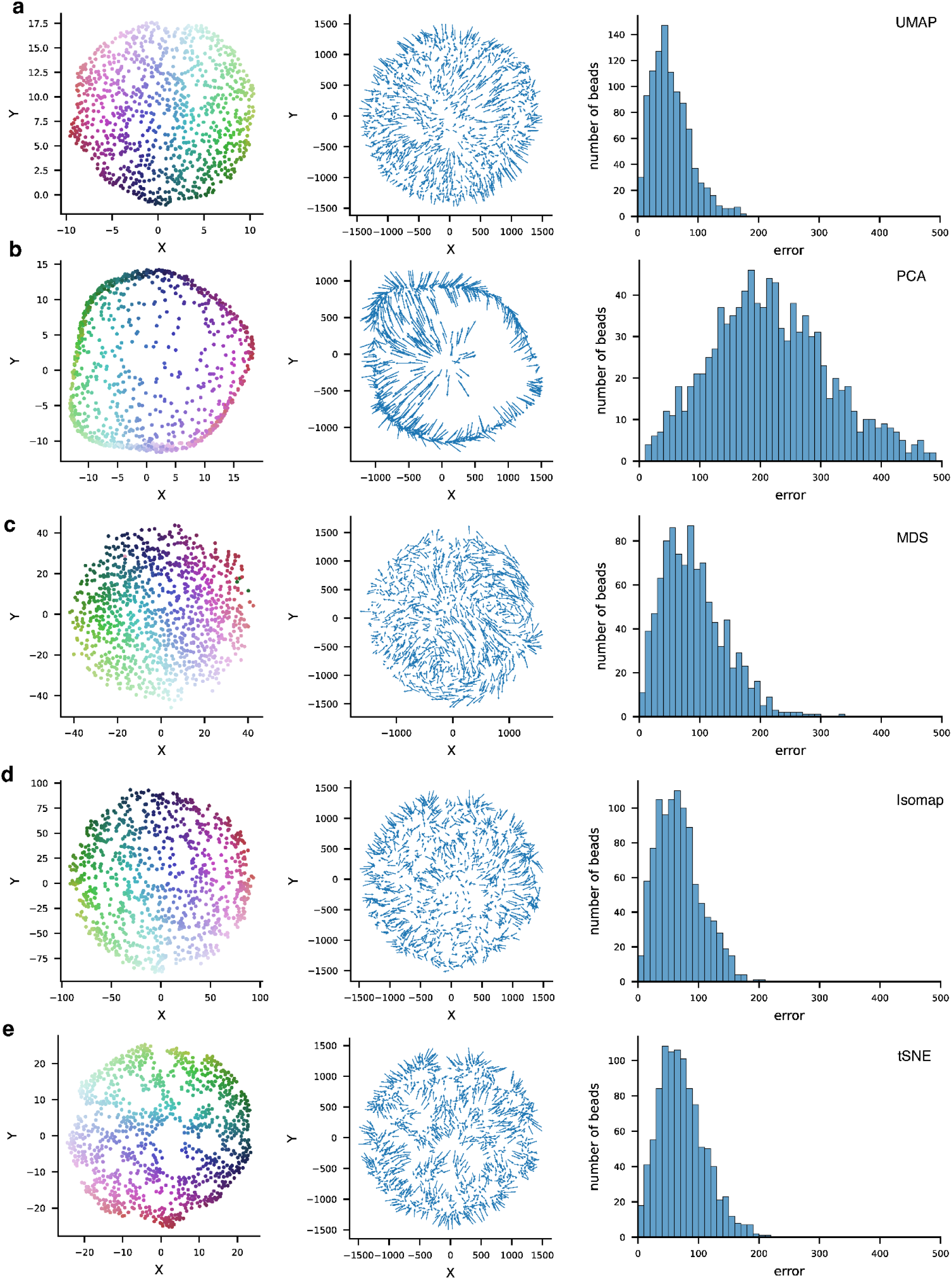
Reconstruction with different dimensionality reduction methods. Reconstruction of identical simulation data with **a**, UMAP, same as Supplementary Fig 1. **b**, Principal component analysis (PCA). **c**, Multidimensional scaling (MDS). **d**, Isomap. **e**, t-distributed stochastic neighbor embedding (t-SNE). Columns: Left, Reconstructed locations of capture beads, colored the same as in Supplemental Figure 1c. Middle, Displacement vectors of capture beads. Each arrow starts from the capture bead’s ground truth location and ends at the reconstruction location. Right, Histogram plot of capture beads’ absolute error.

**Supplemental Figure 4:**
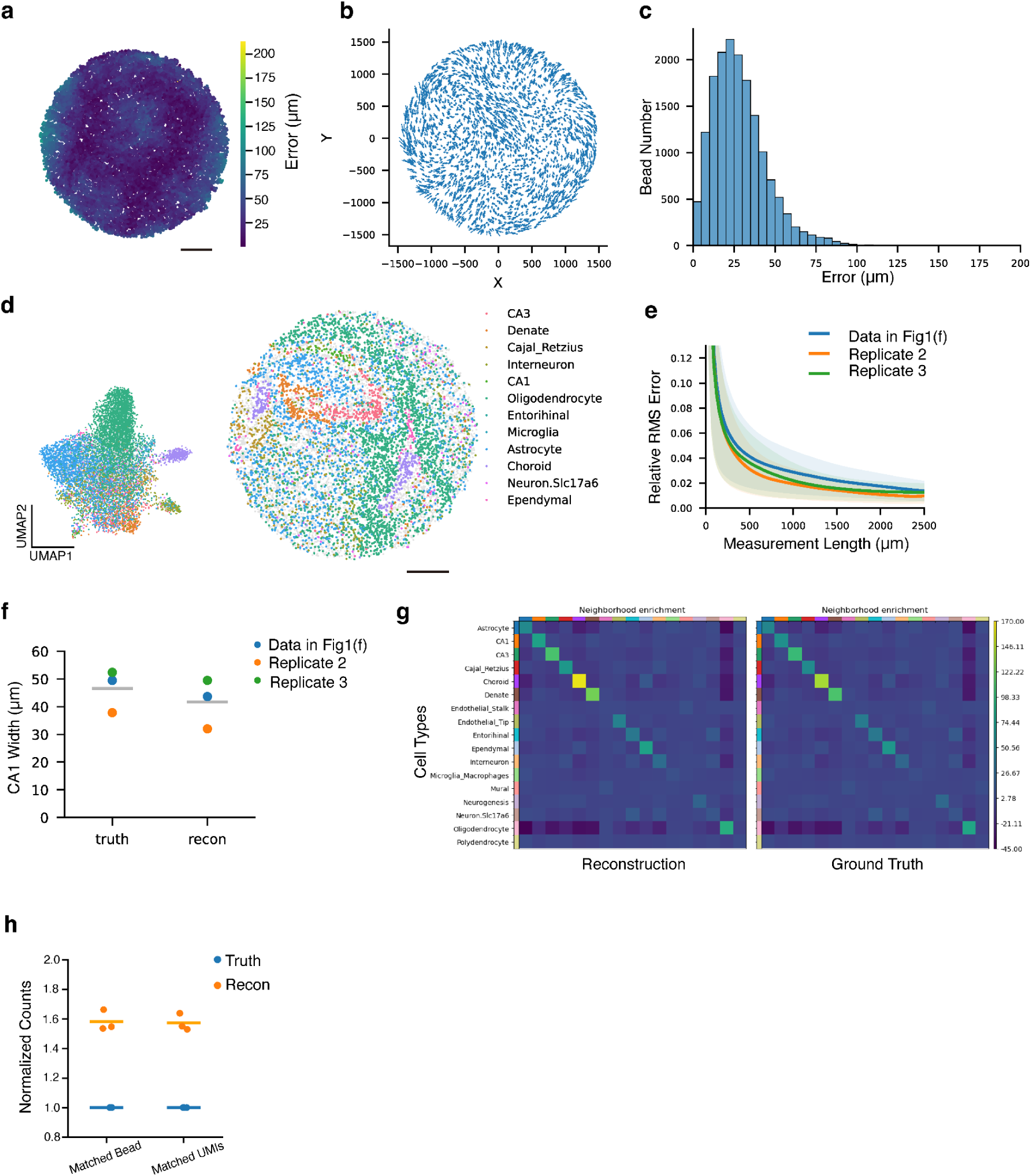
Slide-seq reconstruction metrics. **a**, Absolute error of capture beads plotted in ground truth locations. **b**, Displacement vectors of capture beads. Each arrow starts from the capture bead’s ground truth location and ends at the reconstruction location. **c**, Histogram plot of capture beads’ absolute errors. **d**, (Left) UMAP representing gene expression from a coronal mouse hippocampus section captured by beads, colored by decomposed cell types from RCTD. (Right) Spatial location of capture beads in ground truth, colored by decomposed cell types. **e**, Relative RMS error of measurement lengths as a function of measurement length. Data shown in Fig.1f (blue) and two biological replicates (orange and green) are presented. Solid lines represent average values across beads and shaded areas represent one standard deviation. **f**, CA1 width measured in ground truth and reconstruction (N = 3 biological replicates). Data shown in Fig.1f (blue) and two biological replicates (orange and green) are shown. Gray lines showed the mean width of each group. **g**, Neighborhood enrichment analysis between cell type pairs in reconstruction (left) and ground truth (right). The enrichment scores are plotted in the same color scale, higher scores represent more enriched in the neighborhoods. **h**, Comparison of matched bead barcodes and unique RNA molecules (UMIs) in reconstruction (orange) or ground truth (blue) (N = 3 biological replicates). Solid lines: mean values. Values are normalized to the ground truth for direct comparison across replicates. Scale bars: 500 µm.

**Supplemental Figure 5:**
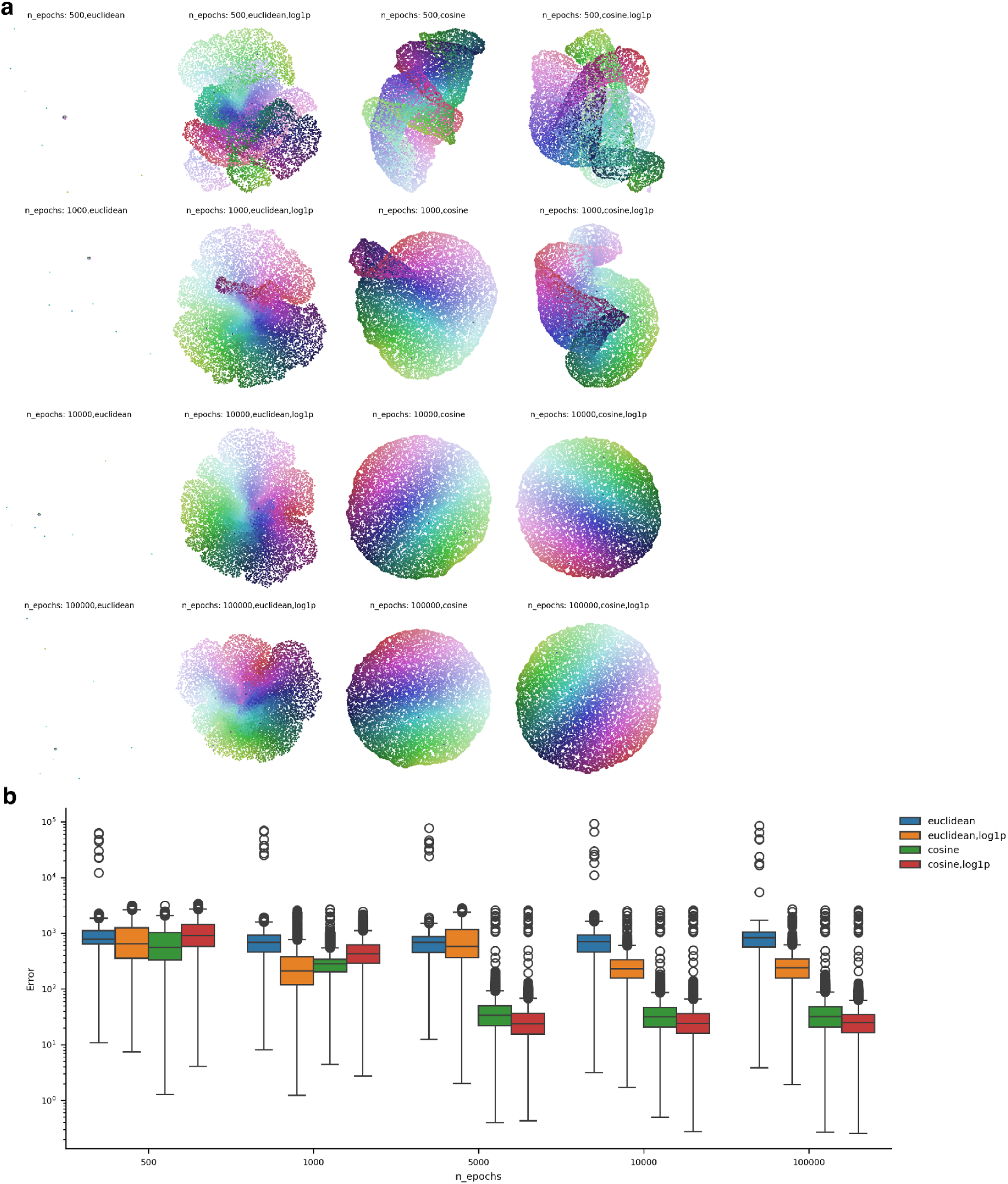
UMAP parameters and matrix normalization effect on reconstruction on experimental data. **a**, The same diffusion matrix is embedded in the two dimensional space with different parameters of UMAP: n_epochs as the number of training epochs and metric as ways of computing distances in high dimensional space^16^. We also tested the effect of applying log1p transformation before UMAP embedding. Each bead is colored according to ground truth location to show the pattern recovery. **b**, Boxplot of reconstruction error with different UMAP parameters and log1p transformation. Each box represents absolute errors of beads with corresponding parameters, with middle box as error between 25% to 75%, middle line as median error, and outlier as 1.5 times of interquartile range (IQR).

**Supplemental Figure 6:**
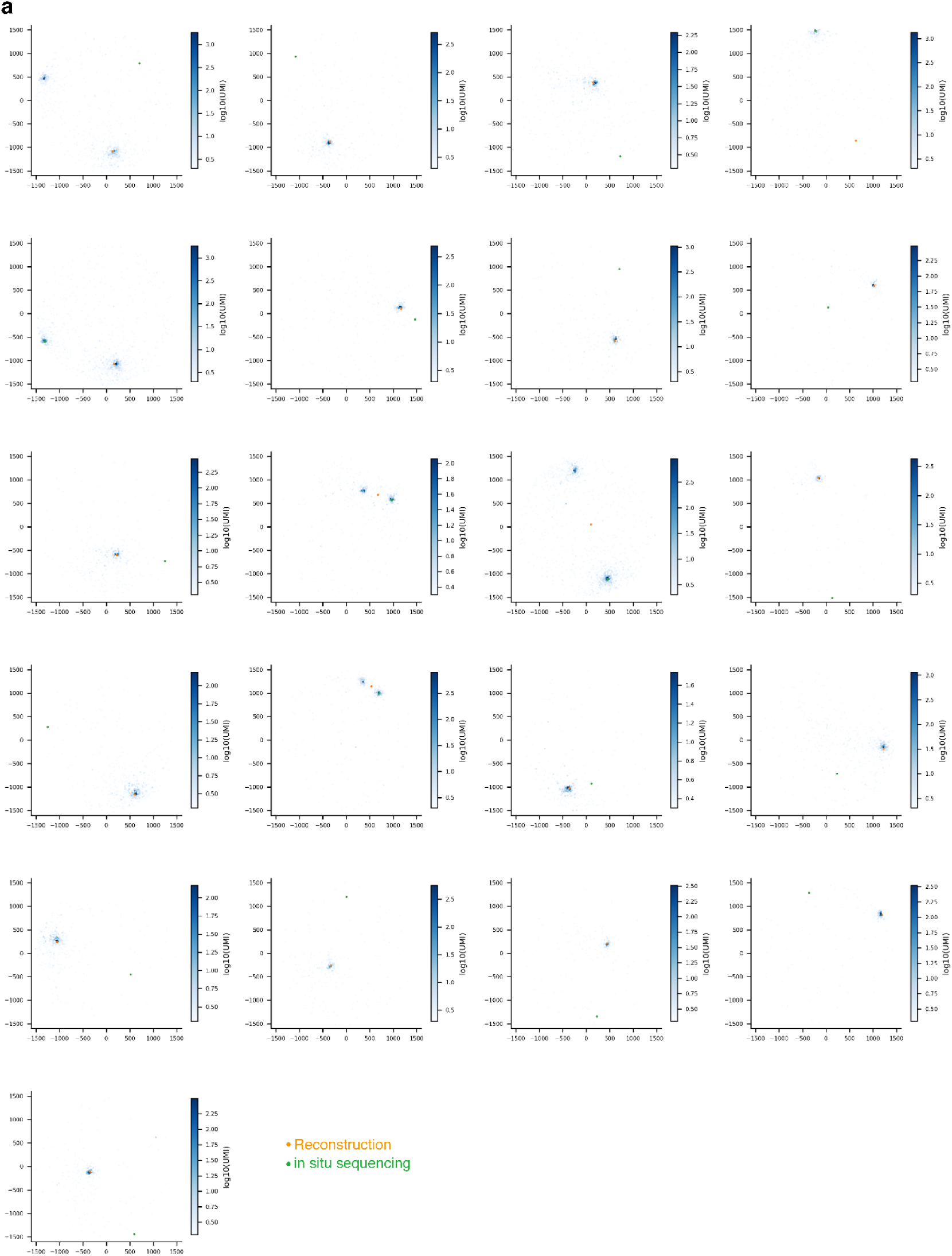
Diffusion and locations of all distant (>200 µm) positioned beads. **a**, 21 capture beads with absolute error >200 µm are displayed with diffusion distributions (blue gradients), reconstruction locations (yellow dots), and in situ sequencing locations (green dots). Among the 21 beads, 5 beads are displayed as doublets (barcode collisions, expected from length of barcode and number of beads), 15 beads have in situ sequencing locations far away from diffusion distribution (we anticipate these to be likely in situ sequencing error, or illumina sequencing error), and one bead has reconstruction location far away from diffusion distribution.

**Supplemental Figure 7:**
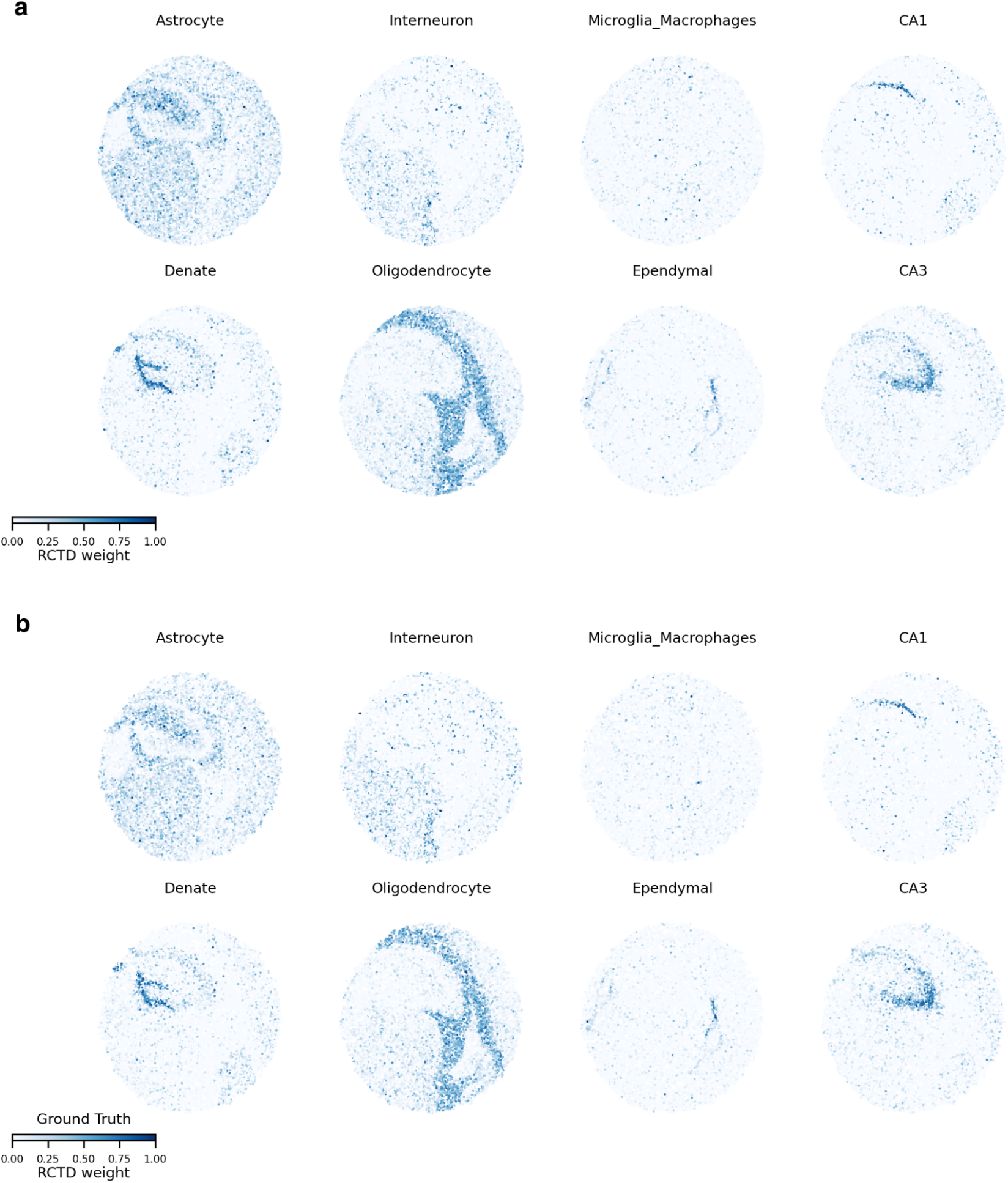
Spatial distribution of cell type RCTD weights in reconstruction and ground truth. Spatial representation of RCTD inferred cell type weights in **a**, Reconstruction. **b**, Ground truth.

**Supplemental Figure 8:**
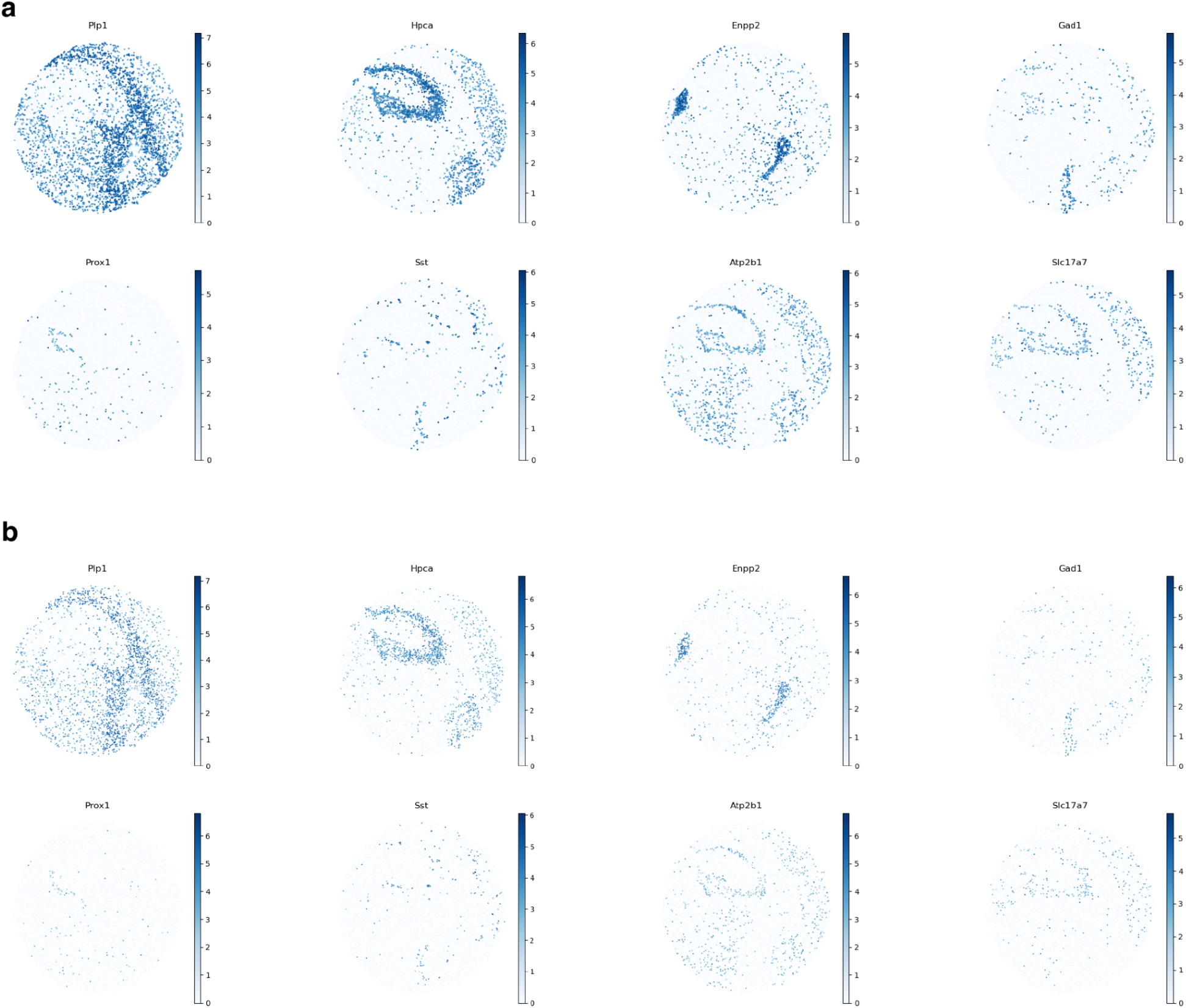
Spatial distribution of genes in reconstruction and ground truth. Spatial expression pattern of marker genes in **a**, Reconstruction. **b**, Ground truth.

**Supplemental Figure 9:**
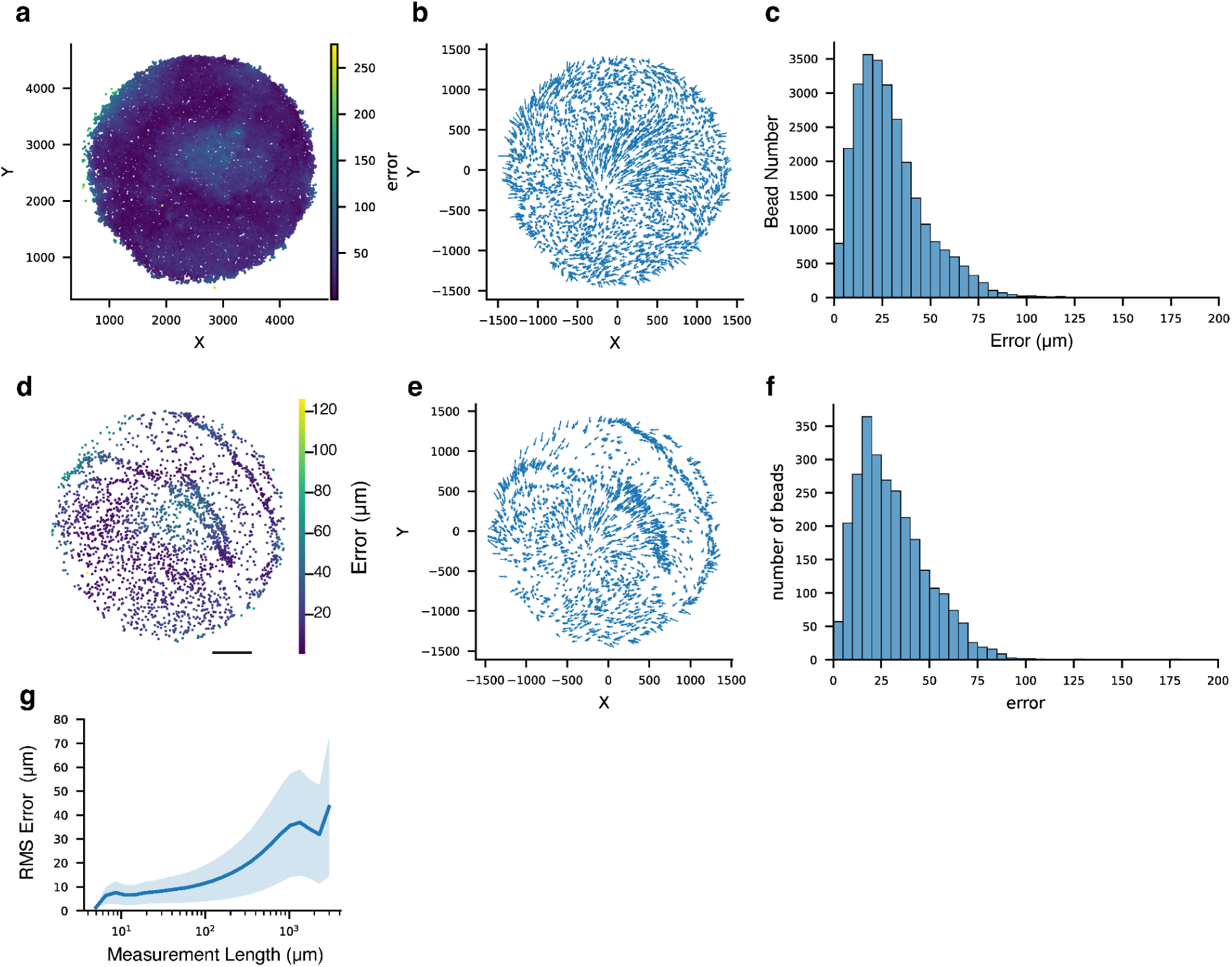
Slide-tags Reconstruction metrics. **a**, Absolute error of capture beads plotted in ground truth locations. **b**, Displacement vectors of capture beads. Each arrow starts from the capture bead’s ground truth location and ends at the reconstruction location. **c**, Histogram plot of capture bead locations’ absolute errors. **d**, Spatial representation of reconstruction error on each nuclei. **e**, Displacement vectors of located nuclei. Each arrow starts from the nuclei’s ground truth location and ends at the reconstruction location. **f**, Histogram plot of nuclei locations’ absolute errors. **f**, RMS error of measurement lengths between bead pairs as a function of measurement length. Solid lines represent average values and shaded areas represent one standard deviation.

**Supplemental Figure 10:**
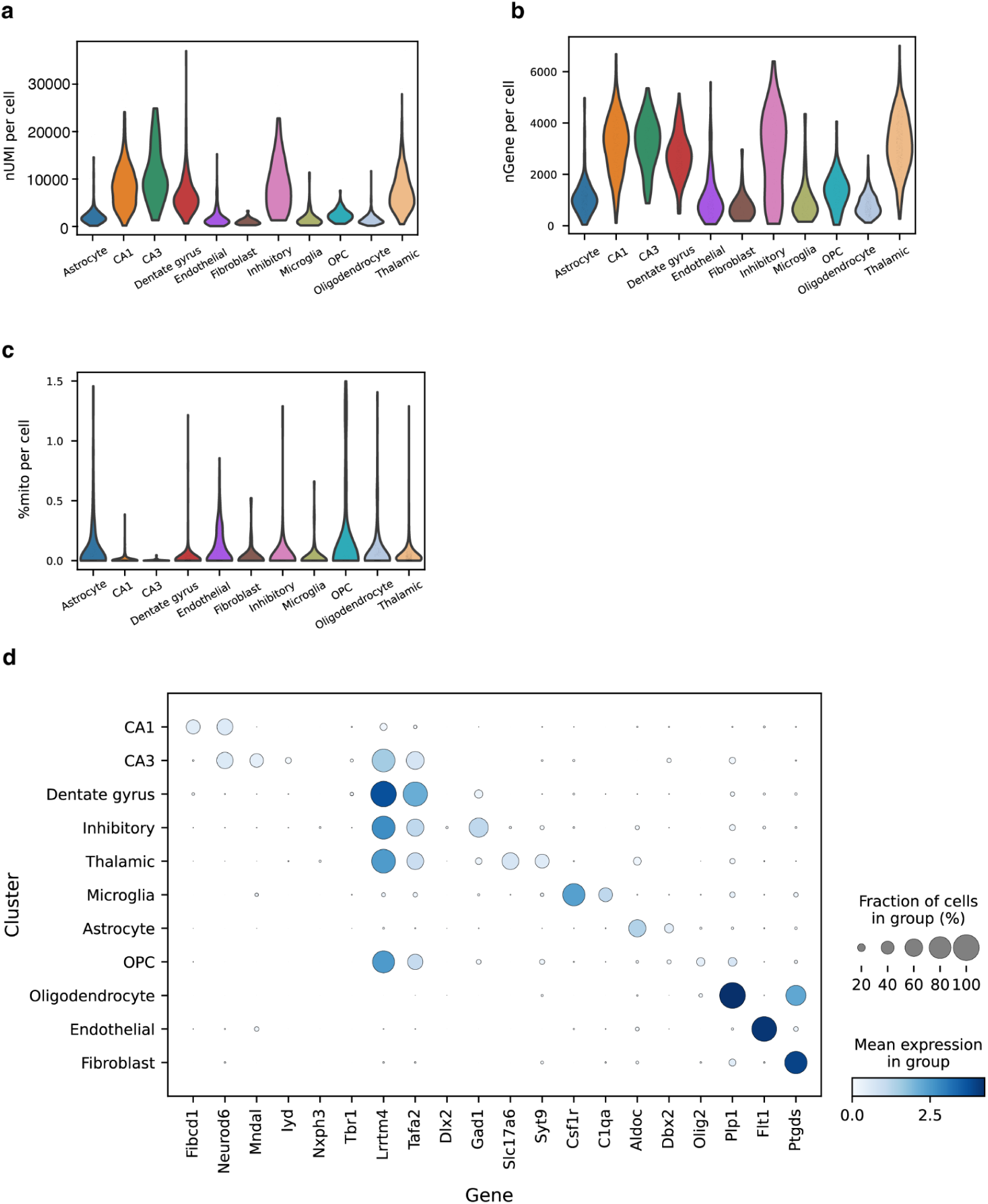
snRNA-seq data metrics for Slide-tags reconstruction. **a**, Distribution of the number of UMI per cell for each cluster. **b**, Distribution of the number of genes per cell for each cluster. **c**, Distribution of the percentage of mitochondrial genes per cell for each cluster. **d**, Expression of marker genes by cell type cluster.

**Supplemental Figure 11:**
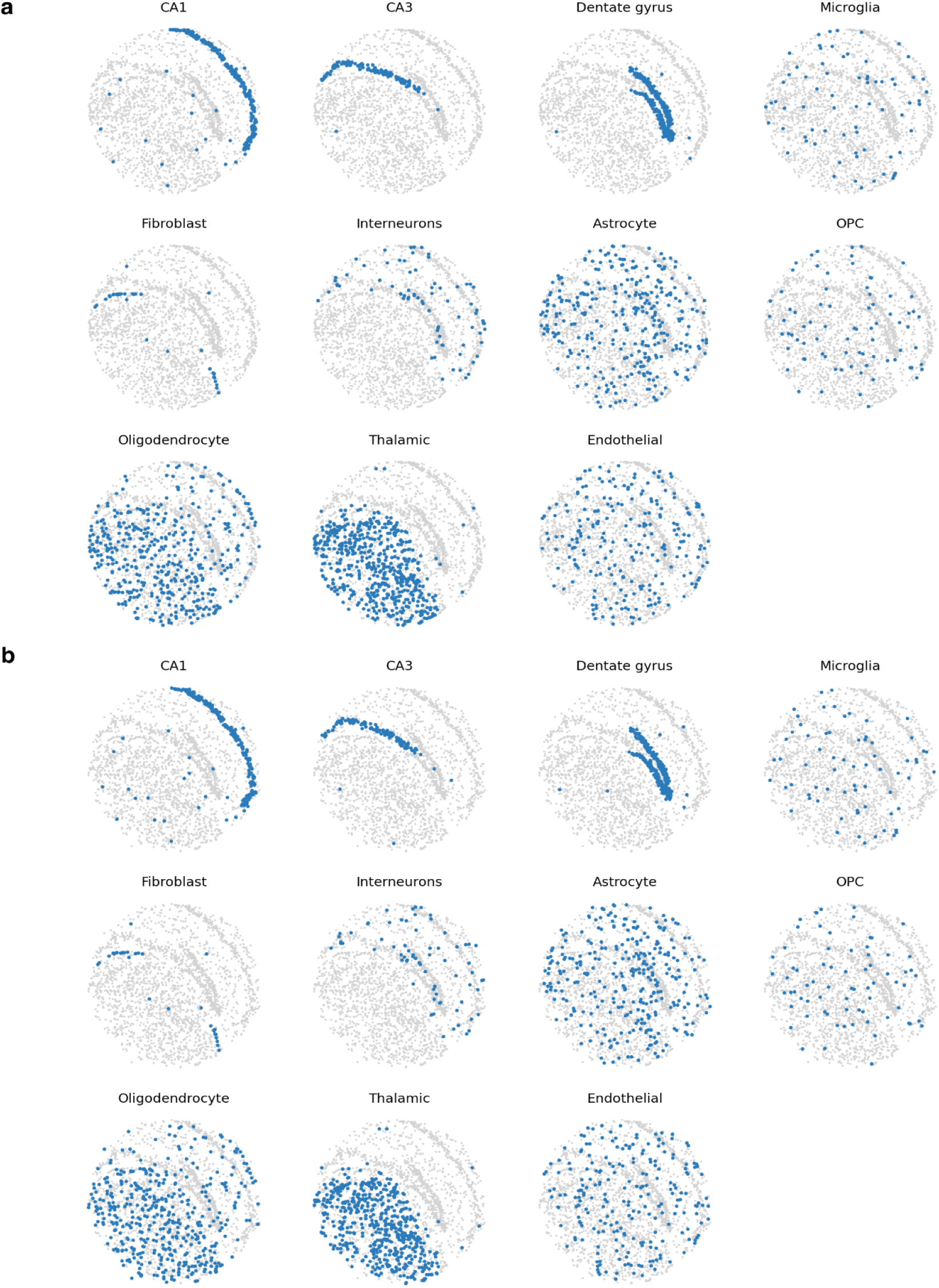
Spatial representation of cell types in reconstruction and ground truth. Spatial distribution of located cell types in **a**, Reconstruction. **b**, Ground truth.

**Supplemental Figure 12:**
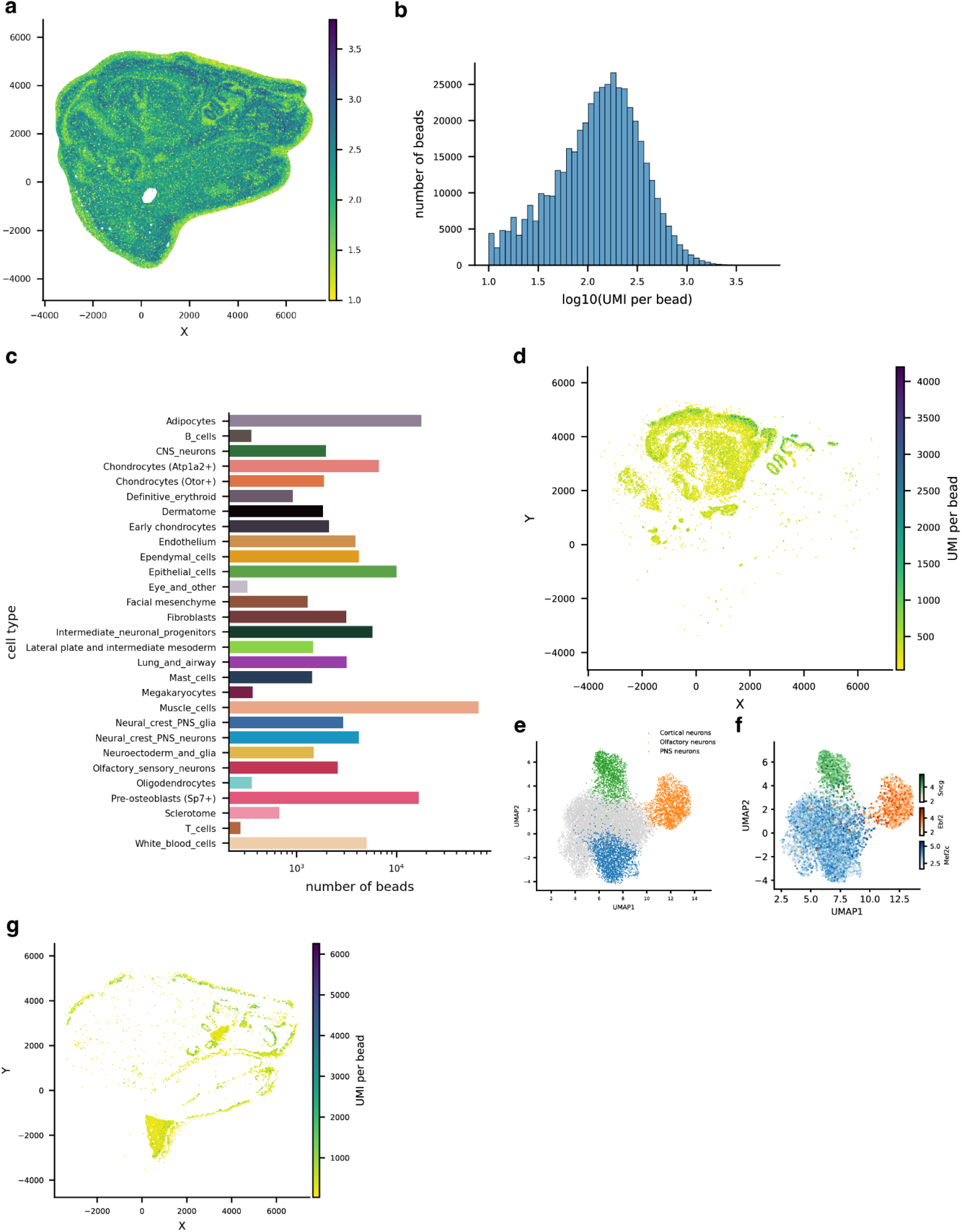
Slide-seq Reconstruction for P1 mouse sample. **a**, Spatial representation of UMI per bead on reconstructed locations. **b**, Distribution of UMI per bead. **c**, Number of beads with assigned cell type from RCTD. d, Spatial representation of UMI per bead of beads assigned as neuronal cells. **e**, UMAP embedding of re-clustered neuronal cells with some cell types labeled. **f**, Marker gene expression of labeled cell types in e on UMAP embedding. **g**, Spatial representation of UMI per bead of beads assigned as epithelial cells.

**Supplemental Figure 13:**
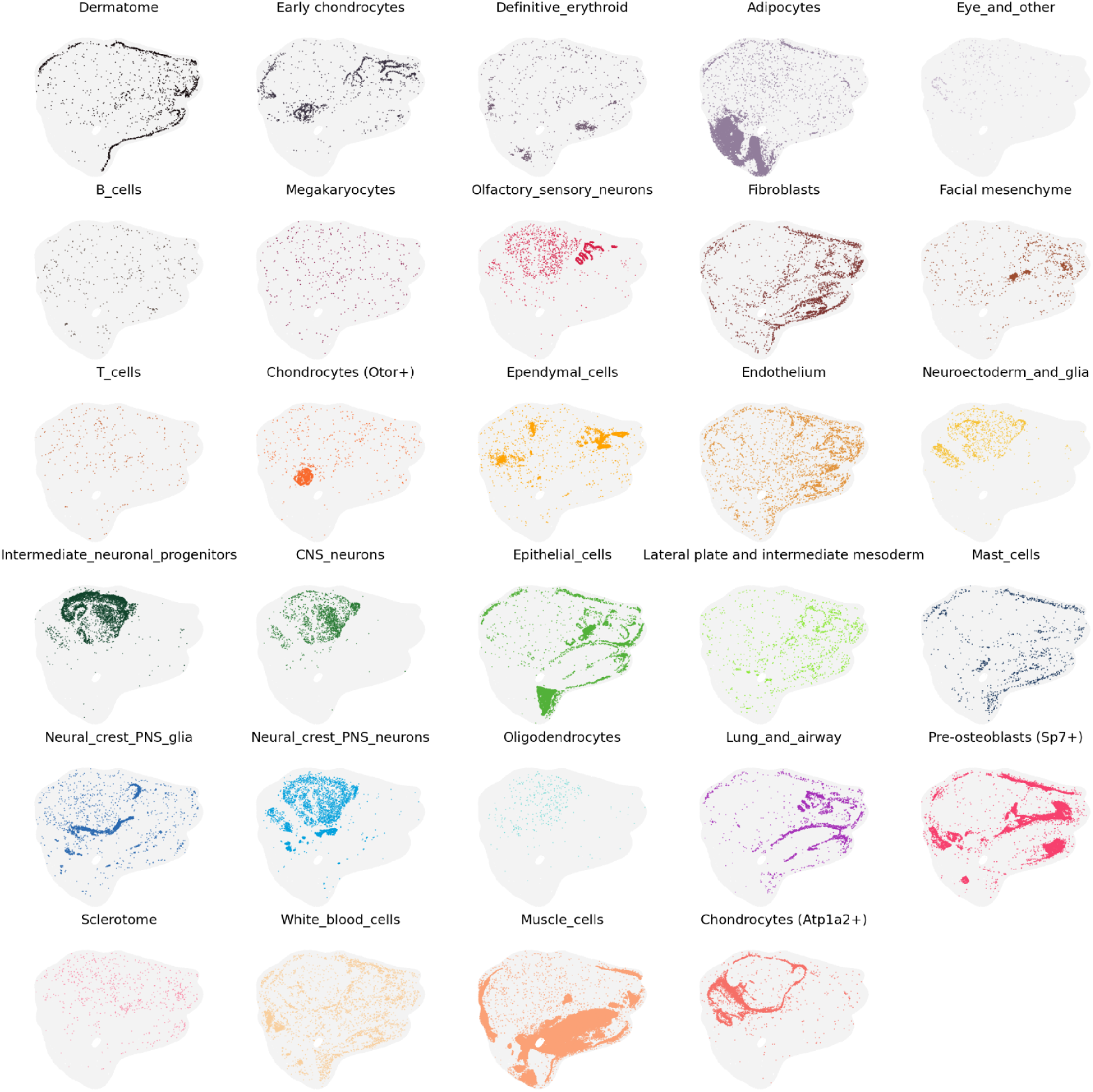
Spatial representation of cell types in reconstruction of P1 mouse sample. Spatial distribution of cell types from RCTD.

**Supplemental Figure 14:**
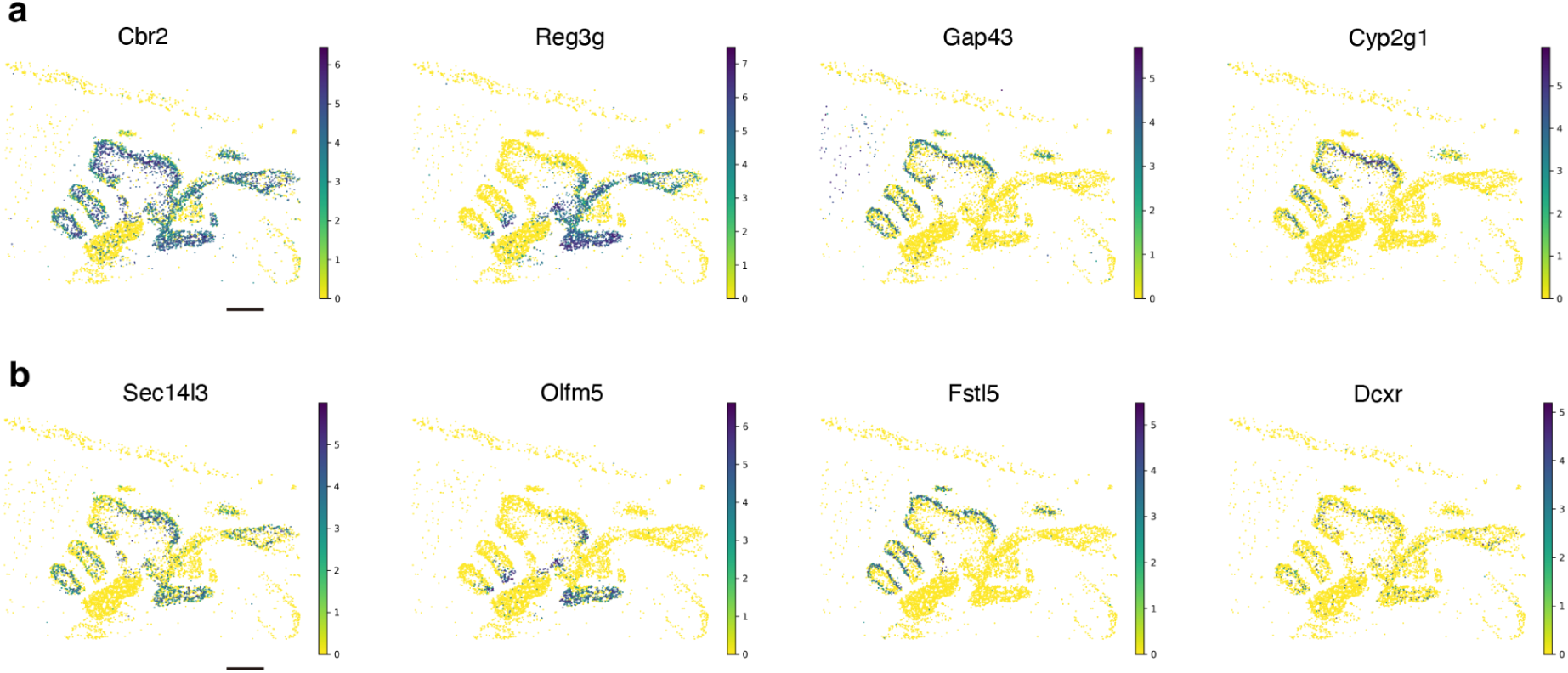
Spatially differential expression genes in olfactory epithelium and respiratory epithelium. **a**, Spatial representation of previously characterized olfactory and respiratory differential expression genes. **b**, Spatial representation of another four less well characterized olfactory and respiratory differential expression genes that were found in this spatial data. Scale bars: 500 µm.

**Supplemental Figure 15:**
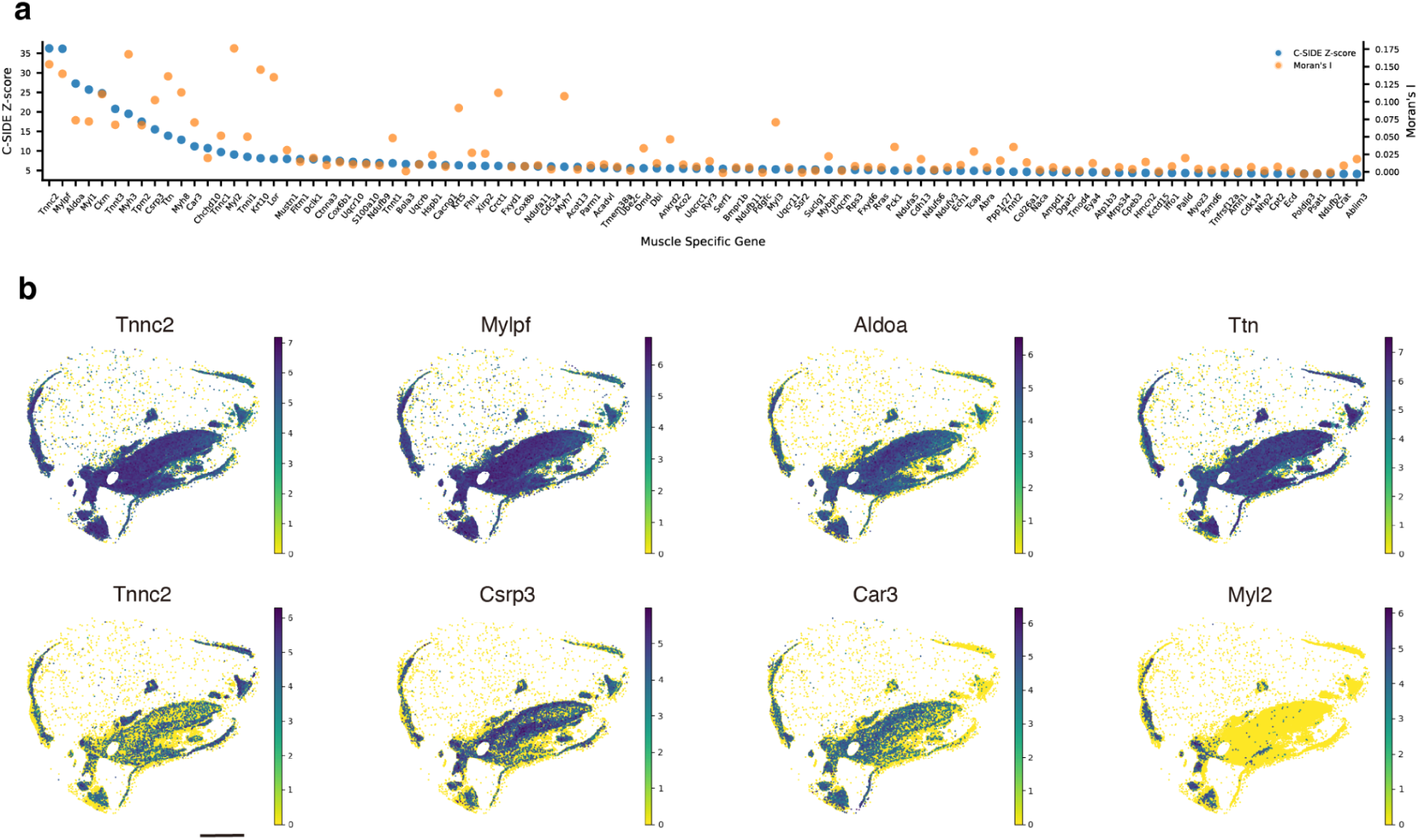
Spatially differential expression genes in muscle cells. **a**, Top 100 spatially differential expression genes of muscle cells were ranked by nonparametric C-SIDE Z-score, with Moran’sI statistics calculated. Higher scores on both metrics signify more spatially variable expression. **b**, Spatial representation of eight spatially differential expression genes of muscle cells from b. All beads of epithelial type are positioned, colored by relative expression level of each gene. Scale bar: 2 mm.

## Supplementary Tables and Videos

Supplementary Video 1: UMAP epochs during reconstruction of P1 Mouse Embryo

Supplementary Table 1: C-SIDE significant differential expression genes

Supplementary Table 2: Reconstruction cost and production time

## Code availability

All code is available at github.com/Chenlei-Hu/Slide_recon.git.

## Data availability

All data is available at the Broad Single Cell Portal: singlecell.broadinstitute.org/single_cell/study/SCP2577. Raw sequencing data will be available on SRA upon publication.

## Acknowledgements

We thank D. Sun, R. Raichur, and S. Alakwe for assistance with tissue handling and library preparation; D. Cable for the initial computational implementation of the idea; K. Cao, J. Zhang, M. Dai, X. Ye, and P. Yadollahpour for discussion on the computational algorithm. Components of the figures were created using BioRender. This work was supported by the National Institutes of Health (grant nos. R01HG010647 to F.C.) and the Opportunity Fund through the Technology Development Coordinating Center at Jackson Laboratories (National Human Genome Research Institute U24HG011735). F.C. also acknowledges support from the Searle Scholars Award, the Burroughs Wellcome Fund CASI award, and the Merkin Institute.

## Competing interests

F.C. is an academic founder of Curio Biosciences and Doppler Biosciences, and scientific advisor for Amber Bio. F.C’s interests were reviewed and managed by the Broad Institute in accordance with their conflict-of-interest policies. E.Z.M. is an academic founder of Curio Biosciences.

## Methods

### Sample information and processing

Mouse brain samples were obtained following guidelines in accordance with the US National Institutes of Health Guide for the Care and Use of Laboratory Animals under protocol number 0120-09-16 and approved by the Broad Institutional Animal Care and Use Committee. Wild-type C57BL/6 mice, maintained on a 12-hour light/dark cycle, were anesthetized by administration of isoflurane in a gas chamber flowing 3% isoflurane for 1 min. Blood was cleared from the brain using transcardial perfusion with a chilled pH 7.4 HEPES buffer (110 mM NaCl, 10 mM HEPES, 25 mM glucose, 75 mM sucrose, 7.5 mM MgCl2, 2.5 mM KCl). Brain was removed, frozen in liquid nitrogen vapor for 3 minutes and stored at −80 °C.

The C57BL/6 mouse embryo at P1 (MF-104-P1-BL) was purchased from Zyagen. The sample was stored at −80 °C before use.

### Histological processing

A 12 µm thick section from the frozen P1 mouse sample was mounted onto a glass plus slide. The Leica ST5010 Autostainer XL (Leica Biosystems) was used for hematoxylin and eosin (H&E) staining. Sections were immersed in xylene, sequentially processed through 100% and 95 % ethanol series, and then stained with hematoxylin. Eosin staining was applied and the section was again processed through 100% and 95 % ethanol series, xylene, dehydrated, and covered using the Leica CV5030 Fully Automated Glass Coverslipper. The slide was imaged with Leica Aperio VERSA Brightfield, Fluorescence & FISH Digital Pathology Scanner.

### Barcoded beads and array production

Bead barcodes were synthesized as in Slide-seqV2^17^. Sequences of beads used in reconstruction for Slide-seq and Slide-tags:

1. Capture beads: 5’-TTTTCTACACGACGCTCTTCCGATCTJJJJJJJJTCTTCAGCGTTCCCGAGAJJJJJJJNNNNN NNVVT30-3’;
2. Fiducial beads: 5’-TTT-pc-GTGACTGGAGTTCAGACGTGTGCTCTTCCGATCTJJJJJJJJTCTTCAGCGTTCCCG AGAJJJJJJJNNNNNNNVVA30-3’, ‘pc’ in the sequence denotes photocleavable linker.
3. Sequence of capture beads used in reconstruction of P1 mouse sample: 5’-PEG-pc-TTTCTACACGACGCTCTTCCGATCTJJJJJJJJTCTTCAGCGTTCCCGAGAJJJJJJJN NNNNNNVVT30, ‘PEG’ in the sequence denotes a polyethylene glycol (PEG) linker.

In Slide-seq experiments, capture beads and fiducial beads were mixed at the ratio of 3:1. In Slide-tags experiments, capture beads and fiducial beads were mixed at the ratio of 1:3. After mixing of the beads, array preparation and in situ sequencing were performed as described previously^17^.

### Reconstruction procedure and library preparation

For reconstruction with Slide-seq, the protocol of Slide-seq V2^17^ was performed until the step of reverse transcription (RT). After RT, the bead array was put on a glass slide in a 10 µL diffusion buffer (2x SSC, 20% formamide). The bead array was then placed under an ultraviolet (365 nm) light source (0.42 mW mm^−2^, Thorlabs, M365LP1-C5, Thorlabs, LEDD1B) for 2 min (Slide-seq for mouse hippocampus section) or 5 s (Slide-seq for P1 mouse sample, as the capture beads used were also photocleavable so we cleaved less time). Then the bead array was incubated at room temperature for 10 min for cleaved oligonucleotides to diffuse. After dipping the bead array into a 1 mL diffusion buffer to wash out free oligonucleotides, the bead array was put into a 1.5 mL centrifuge tube of 200 µL extension buffer (1x NEBuffer 2, 1mM dNTP, 25 units Klenow exo- (NEB M0212L)). The bead array was incubated at 37 °C for 1 h. Then Slide-seq V2 was continued from adding the tissue clearing buffer to cDNA PCR on the dissociated beads. After cDNA PCR, the beads were spinned down with 2 min 3000 RCF. Then supernatant was collected for cDNA purification and beads were resuspended in PCR mix again for the reconstruction library. The PCR mix included: 1x KAPA (Roche KK2612), 100 nM P5-Truseq Read 1 primer AATGATACGGCGACCACCGAGATCTACACTCTTTCCCTACACGACGCTCTTCCGATCT, 100 nM P7-Truseq Read 2 primer CAAGCAGAAGACGGCATACGAGANNNNNNNNGGTGACTGGAGTTCAGACGTGTGCTCTTC CGATCT.

The reconstruction library PCR followed the program: 3 min at 95 °C, 13 cycles of (20 s at 98 °C, 15 s at 65 °C, 15 s at 72 °C), and 1 min at 72 °C. After the PCR program, supernatant was collected after centrifuge and purified with 1.5x SPRI cleanup (Beckman Coulter, A63881), generating reconstruction library ready for sequencing. Meanwhile, the mRNA library was prepared as described previously^17^.

For reconstruction with Slide-tags, the diffusion and extension steps were performed before doing Slide-tags. Similar to the process in reconstruction with Slide-seq, the bead array was emerged with diffusion buffer and exposed to ultraviolet (365 nm) light source for 5 s, incubated at room temperature for 10 min, dipped in 1 mL diffusion buffer, put in 200 µL extension buffer as above and incubated at 37 °C for 1 h. After extension, the bead array was dried, and used for Slide-tags protocol until nuclei extraction. Afterwards, extracted nuclei followed Slide-tags protocol^19^ for gene expression while beads were dissociated, washed with 200 µL wash buffer (10 mM Tris pH 8.0, 1 mM EDTA and 0.01% Tween-20), and resuspended in 200 µL PCR mix. PCR mix and program are both the same as in reconstruction with Slide-seq. Reconstruction library was ready for sequencing after 1.5x SPRI cleanup.

### Sequencing

The gene expression libraries and reconstruction libraries were sequenced on the Illumina Nextseq1000 instrument using P2 100 cycle kit (Illumina, 20046811). For the P1 mouse sample, the gene expression library was sequenced on an Illumina NovaSeq instrument using the S Prime platform.

### Diffusion matrix simulation

Fiducial and capture beads were simulated to locate uniformly in a circle with each bead’s color determined by its location in the image of pattern ‘H’ or in a two dimensional color gradient. The diffusion of a fixed number of barcodes, described by unique molecular identifiers (UMIs), from each fiducial bead was assumed to follow a Gaussian distribution. *Y_ij_*, the number of fiducial bead barcode *j* captured by capture bead *i*, follows a binomial distribution with the probability determined by the distance between beads

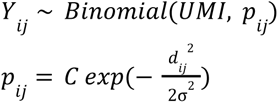

where *d_ij_* is the Euclidean distance between the fiducial and capture beads, σ is the standard deviation in the Gaussian distribution, and *C* is for normalization. The diffusion based pairwise count matrix was then generated for reconstruction.

For the simulated diffusion matrix, 1000 capture beads and 1000 fiducial beads were located uniformly with a circle of diameter as 3000 µm. Diffusion distance σ was 300 µm and UMI was 300 when simulating the diffusion matrix.

### Sequencing data processing and diffusion matrix generation

In the processing of FASTQ files from the reconstruction library sequencing, reads were initially filtered out if their constant sequences had Hamming distances greater than 3 when compared to the universal primer sequence. From remaining reads, capture bead barcode, capture bead UMI, fiducial bead barcode, and fiducial bead UMI were abstracted. To determine the read threshold for reliable bead barcodes, rank plots were generated for both capture bead barcodes and fiducial bead barcodes. Barcodes above the read threshold were collapsed with a Hamming distance of 1, resulting in a whitelist of barcodes. Barcodes with reads below the threshold were matched to the whitelist with a Hamming distance of 1. Paired reads with both capture bead barcode and fiducial bead barcode in the whitelist were compiled along with the UMI information.

Data frames of capture bead barcode, fiducial bead barcode, and UMI were grouped by the combinations of capture bead barcode and fiducial bead barcode. Within these groups, the unique UMIs were counted, with these counts serving as elements within diffusion matrices. Diffusion matrices in sparse matrix data structure were generated by taking capture bead barcodes as rows and fiducial bead barcodes as columns (or fiducial bead barcodes as rows and capture bead barcodes as columns for reconstruction with Slide-tags). The element values in these matrices corresponded to the count of unique UMIs associated with each barcode pair.

### Computational reconstruction with UMAP

Diffusion matrices were used as input for Uniform Manifold Approximation and Projection^16^ (UMAP) to reduce to a two dimensional space. Coordinates in the two dimensional space were directly used as reconstructed locations. UAMP parameters that we tuned are: larger n_neighbors and larger min_dist for uniform distribution of beads, larger n_epochs for converging, and also cosine metric that can better represent the high dimensional distance from diffusion matrix. With experimental data, we also found a log1p transformation of the diffusion matrix improved reconstruction accuracy.

For simulated diffusion matrix, we used UMAP to embed the diffusion matrix directly in a two-dimensional space with the following parameters: cosine metric, n_neighbores=25, min_dist=0.99, n_epochs=10000, learning_rate=1.

For reconstruction with Slide-seq or Slide-tags, we used UMAP to embed the log1p transformed diffusion matrix in a two-dimensional space with the following parameters: cosine metric, n_neighbores=25, min_dist=0.99, n_epochs=50000, learning_rate=1. The UMAP computation was expedited through the use of 24 parallel threads.

For reconstruction with P1 mouse, we used UMAP to embed the log1p transformed diffusion matrix in a two-dimensional space with the following parameters: cosine metric, n_neighbores=45, min_dist=0.4, n_epochs=10000, learning_rate=1. The UMAP computation was expedited through the use of 24 parallel threads. We changed the parameters for this 1.2 cm sample because: the diffusion distance (σ) was the same while the whole size increased, which means the relative connectivity decreased and the minimum distance between beads decreased.

### Comparing reconstruction with other dimensionality reduction methods in simulation

The same simulated diffusion matrix was used for testing and comparing other dimensionality reduction methods.

PCA was performed with n_components=2 and other default parameters; MDS was performed with n_components=2 and other default parameters; Isomap was performed with n_components=2, n_neighbors=100, max_iter=100000; tSNE was performed with n_components=2, random_state=0, perplexity=50.

### Evaluation of reconstruction results

To estimate the absolute errors of beads locations, reconstructed locations were registered to ground truth (from simulation or in situ sequencing) using procrustes analysis^23^, which applies only rigid transformation (scaling, reflection, rotation, translation) on the reconstructed locations. The reconstruction error of each bead was calculated directly by comparing the locations in ground truth and in transformed reconstruction results. The absolute errors were presented through spatial distributions, displacement vectors, and distribution histograms. In the displacement vectors, only vectors with a length of less than 282.8 µm were included. In distribution histograms, errors were displayed within the indicated ranges.

To estimate the measurement length errors, pairwise distances between capture beads were calculated in both reconstruction and ground truth. The differences between the distances in reconstruction and ground truth were calculated as the measurement length errors. The measurement lengths were categorized into bins with a width of 30 µm, and the errors within each bin were averaged using the root mean square (RMS) method. The standard deviation for the measurement length errors within each bin was also determined. To obtain relative errors, the RMS errors were divided by the corresponding measurement lengths.

### Analysis of diffusion in ground truth

The diffusion distribution of a capture bead barcode was represented by the position of its associated fiducial bead barcodes, which were color-coded according to the conjugation UMI count. The diffusion distributions of capture beads with high total UMI counts (first 3000 beads ranked by total UMI counts) along the X axis were fitted using Kernel Density Estimation (KDE) and then averaged to derive the ensemble KDE diffusion distribution. The empirical Full Width at Half Maximum (FWHM) of the diffusion distribution was calculated based on the ensemble KDE.

### Slide-seq gene expression data analysis

After sequencing, gene expression results were processed using the Slide-seq pipeline without bead barcode matching. We visualized the gene expression data of the beads using UMAP with top 40 principal components and the number of nearest neighbors =10. Then we applied robust cell type decomposition^18^ (RCTD) to decompose the cell type associated with each capture bead barcode.The single-cell reference dataset used for comparison was the same as the one for the mouse hippocampus mentioned in the RCTD study^17^.

### CA1 width analysis

We profiled the spatial distribution of Atp2b1in both reconstruction and ground truth. A line perpendicular to the expression pattern of Atp2b1 in the CA1 region was drawn to characterize the expression density of Atp2b1 along this line. The widths of the CA1 region were then determined by the Full Width at Half Maximum (FWHM) of the Atp2b1 expression distribution. This analysis of the CA1 width was conducted across three biological replicates.

### Neighborhood enrichment analysis

The neighborhood enrichment analysis was performed using the squidpy^24^ package in python (v.1.2.2). Cell types were defined by RCTD and locations were from reconstruction and ground truth respectively. Results from reconstruction and ground truth were presented in the same scale. Pearson correlation^23^ was calculated for the z-score of neighborhood enrichment from reconstruction and ground truth, with a ‘two-sided’ alternative hypothesis.

### Slide-tags gene expression data analysis and nuclei positioning

In reconstruction with Slide-tags, single-nucleus gene expression data and spatial barcode data were analyzed according to the Slide-tags pipeline. Nuclei positioning was performed using bead locations from both the reconstructed data and the ground truth respectively.

### Evaluation of nuclei positioning error

To evaluate the error in nuclei positioning, we first transformed nuclei locations from reconstruction according to the registration between bead locations in reconstruction and ground truth. We compared nuclei locations from the reconstruction and the ground truth directly to calculate the absolute error. For the analysis of the measurement length error, we examined the pairwise distances between nuclei, employing the same methodology used for the measurement length error of beads.

### Segmentation of P1 mouse bead array

A 1.2 cm circular bead array was used to profile the spatial transcriptomics of the P1 mouse section although the tissue covered only a portion of the bead array. To differentiate between the tissue-covered and uncovered regions of the bead array, segmentation was performed based on the UMI (Unique Molecular Identifier) count per bead. Kernel Density Estimation (KDE) was employed to estimate the UMI count density across the array, and a threshold for UMI counts was established. Only the beads covered by tissue were retained for further analysis, to save computational memory.

### P1 mouse gene expression analysis

After sequencing, the gene expression results were processed using the Slide-seq pipeline. The gene expression data of the beads were visualized using UMAP, with top 40 principal components and the number of nearest neighbors =10. Then we applied RCTD to decompose the cell type associated with each capture bead barcode with a reference from P1 mouse single cell data^25^. In the reference, we excluded cell types that were not present in the tissue section being profiled.

Neuronal cell types, including CNS neurons, neural crest and PNS neurons, olfactory sensory neurons, and intermediate neuronal progenitors, were isolated and analyzed again with UMAP embedding and unsupervised clustering. Highly variable genes were found for each subcluster. Olfactory epithelium enriched genes were listed by calculating the ratio of the mean expression level in the olfactory epithelium region to that in the entire section.

To identify genes with spatial differential expression, we performed nonparametric cell type-specific inference of differential expression^21^ (C-SIDE) analysis focusing on epithelial and muscle cells. Furthermore, we calculated Moran’s I statistics^24^ to verify the patterned gene expression detected through C-SIDE.

## Notes

### Summary of Updates

We updated the acknowledgments for a complete list of funding source.

