## Supplementary figures and images for "Scalable imaging-free spatial genomics through computational reconstruction"

### Supplementary Video 1

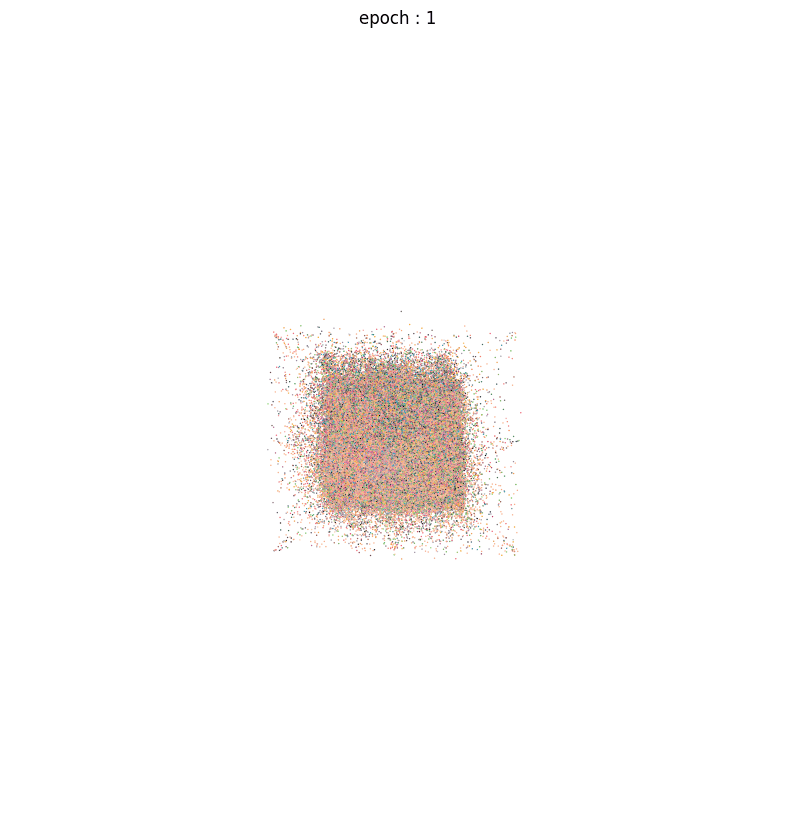
